# Enhanced Detection of RNA Modifications in *Escherichia coli* Utilizing Nanopore RNA004 Technology

**DOI:** 10.1101/2025.01.16.633309

**Authors:** Zhihao Guo, Yanwen Shao, Lu Tan, Beifang Lu, Xin Deng, Sheng Chen, Runsheng Li

## Abstract

RNA modifications are critical regulators of diverse cellular processes, yet their roles in prokaryotic mRNAs remain poorly understood. Recent advances in Oxford Nanopore sequencing—especially the RNA004 kit—have enabled higher yields, reduced signal-to-noise ratio, and improved read accuracy, making them promising tools for investigating bacterial epitranscriptomes. Here, we presented a comprehensive walkthrough for *Escherichia coli* RNA modification analysis based on RNA004. Using both native (WT) and in vitro-transcribed (IVT) RNA samples, we first evaluated the Dorado modification detection models (Ψ, m⁶A, m⁵C, and A-to-I). While each model successfully identified known rRNA modification sites, it also generated many false positives, emphasizing the need for careful data interpretation.

To address these limitations, we introduced nanoSundial (https://github.com/lrslab/nanoSundial), a new comparative method that leveraged raw current features from WT and IVT samples to detect multiple types of RNA modifications in prokaryotes. We optimized nanoSundial on well-studied rRNA sites and validated its effectiveness with tRNA modifications. Through technical and biological replicate analyses, nanoSundial demonstrated reproducibility exceeding 95% in tRNA, rRNA, and ncRNA regions, albeit with lower reproducibility (∼61%) in mRNA. We further found enrichment of mRNA modifications at the start or end of coding sequences. In total, 190 stably modified CDS regions were identified in *E. coli*, many of which cluster near the end of highly expressed transcriptional units (TUs) in each operon.

Overall, this study highlighted the strengths and limitations of current nanopore-based modification detection methods on bacterial RNA, introduced a robust new comparative tool, and elucidated previously uncharacterized mRNA modification landscapes. Our findings open new avenues for understanding the functional impacts of bacterial RNA modifications and advancing epitranscriptomic research in prokaryotes.

## Introduction

Chemical modifications of RNA occur naturally across all domains of life, with over 160 distinct types identified (1–6). However, studying mRNA modifications in bacteria remains especially challenging. For instance, tRNAs and rRNAs are abundant and stable (7), making their modifications relatively easy to study in both eukaryotes (8–10) and prokaryotes (6, 11–13). In contrast, bacterial mRNAs, which lack a poly(A)-tail (14) and have exceptionally short half-lives (15), are more difficult to isolate and characterize. This has made bacterial mRNA modifications particularly challenging to study, even though they likely play significant roles, leaving their modification landscape largely unexplored (16). Recent improvements in bacterial RNA sample preparation allowed a higher mRNA yield and sequencing throughput, facilitating the analysis of bacterial transcriptome and epi-transcriptome (17, 18).

Current RNA modification detection methods, such as liquid chromatography-mass spectrometry (LC-MS) (19–21) and immunoprecipitation-based techniques like MeRIP-Seq (22), have limitations in terms of single-nucleotide resolution (23), target specificity (24, 25), and single-molecule detection (26). Nanopore-based direct RNA sequencing (DRS) offers a promising solution to these challenges (27). RNA modifications generate distinct ionic current signals when RNA molecules pass through nanopores, enabling their detection via DRS (27). Multiple computational tools are developed based on ONT RNA002 kits and can be classified into two categories based on whether their operating characteristics need a control sample. Single-mode methods include tools such as Epinano_SVM (28), nanom6A (29), MINES (30), Tombo_de_novo (31) and m6Anet (26). And comparative methods encompass Epinano_Err (28), nanocompore (32), xPore (33), ELIGOS (34), Tombo_comp (31), DirrErr (35) and DRUMMER (36). However, it is noticeable that these tools could generate highly inconsistent predictions (17, 37).

Recently, Oxford Nanopore Technologies (ONT) introduced an updated Direct RNA Sequencing (DRS) kit (SQK-RNA004) and a corresponding flow cell (FLO-MIN004RA), which deliver higher sequencing yields, reduced current signal noise, and improved quality scores (38). While this advancement opens new opportunities for studying RNA modifications, existing methods optimized for RNA002 cannot be directly applied to RNA004 sequencing data. To address this, ONT officially released the Dorado rna004_130bps_sup@v5.0.0 RNA basecalling model, including several single-mode “all-context” models to detect RNA modifications with RNA004 data. These models had been trained to identify N⁶-methyladenosine (m⁶A) (10, 39–41), pseudouridine (Ψ) (5), Adenosine-to-Inosine (A-to-I) (42), and 5-methylcytosine (m⁵C) (43). Although these models have proven effective in various eukaryotic systems (44), their performance in detecting bacterial RNA modifications remains unverified. Moreover, because previous nanopore-based approaches have shown limited success in detecting DNA modifications in bacteria (45), further validation and exploration of RNA modifications in bacterial RNA004 data are crucial.

Here, we presented the first high-quality bacterial transcriptome data generated via ONT DRS using the RNA004 technology, including wild-type (WT) and *in vitro* transcribed (IVT; unmodified) samples. We were also the first to evaluate the performance of Dorado RNA modification models on bacterial RNA004 data. Furthermore, we introduced nanoSundial, the first de novo comparative approach that utilized current features from WT and IVT RNA for accurate detection of prokaryotic RNA modifications based on RNA004 data. nanoSundial effectively identified multiple types of modifications and was validated on various RNA types, including tRNA, rRNA, and mRNA. Its reproducibility was confirmed through technical and biological replicates. Notably, we observed that mRNA modifications often occur near the start or end of coding sequences (CDS). Overall, this study not only demonstrated the feasibility of identifying bacterial RNA modifications via DRS with RNA004 data, but also highlighted exciting new avenues for investigating the bacterial epitranscriptome.

## Results

### The new RNA004 kit improved sequencing yield and quality for bacterial RNA

Total RNA was extracted from *E. coli* K-12 cells harvested at logarithmic growth phase. Following our previous pipeline, we performed size selection, rRNA depletion, and polyadenylation (ss&rd_RNA) before the DRS library preparation (17). *In vitro* transcribed RNA (IVT_neg) sample was prepared using the ss&rd_RNA as a template (described in Methods section). Sequencing libraries were constructed using the latest SQK-RNA004 kit and ran on the FLO-MIN004RA flowcell. The raw data were basecalled using the Dorado rna004_130bps_sup@v5.0.0 model. We refer to the native samples as ss&rd_004, and the IVT samples as IVT_neg_004.

The ss&rd_004 and IVT_neg_004 samples yielded 6.84 million reads (4201 Mb in bases) and 8.32 million reads (5659 Mb in bases), respectively, representing a more than 4-fold increase compared with previous SQK-RNA002 data (**Figure 1A**). Unfiltered reads were subsequently mapped to the *E. coli* K-12 genome using minimap2. As a result, 93.65% of ss&rd_004 and 94.87% of IVT_neg_004 reads were mapped, corresponding to 92.06% and 90.78% of bases, respectively. We also observed higher read accuracy. The median Q scores were 17.93 for ss&rd_004 and 19.17 for IVT_neg_004 (**Figure 1B, C**), whereas the median Q score with SQK-RNA002 was about 13.

**Figure 1.**
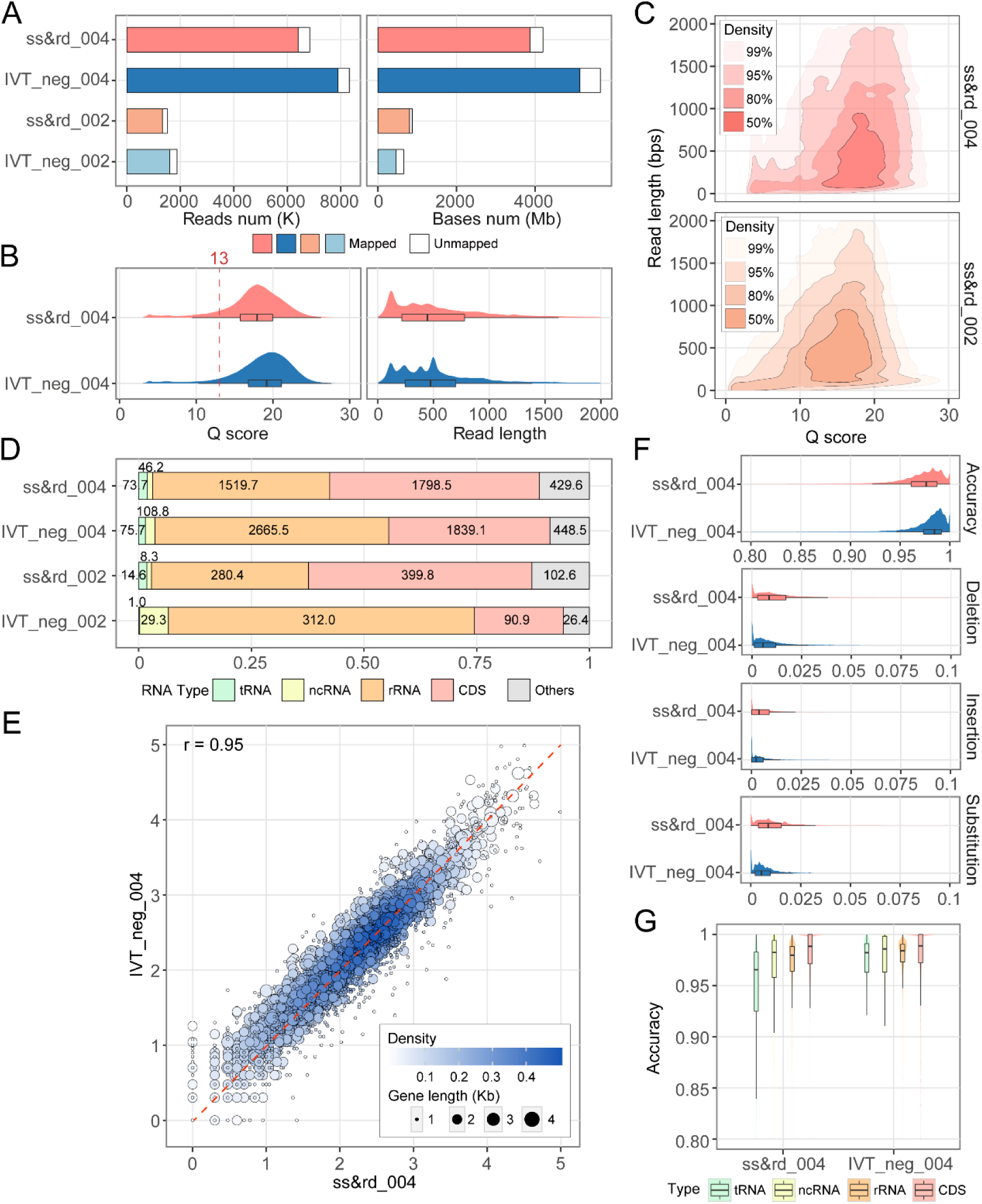
Raw read features and analysis of mapped reads based on Dorado basecalling results. (**A**) Total sequencing throughput from the samples of ss&rd and IVT_neg, sequenced with RNA002 (_002) and RNA004 (_004) technology. ss&rd meant the sample was processed through <u>s</u>ize <u>s</u>election, rRNA <u>d</u>epletion, and polyadenylation. IVT_neg meant the *in vitro*-transcribed sample as a no-modified negative control. The white blank represents the reads and bases cannot align on the reference genome. **(B)** Distribution of Q score and read length from RNA004 raw reads, and 13 is the median Q score of RNA002 data. (**C**) The relationship between read length and quality of raw reads was shown as a 2D density plot. Samples of ss&rd sequenced with RNA004 and RNA002 techniques were displayed. Read qualities were specified by Q scores. (**D**) Proportion of mapped bases to annotation features, including tRNA, ncRNA, rRNA CDS and others. Numbers embedded in the bars indicated the number of bases in each category. Others covered the UTR region and intergenic region. (**E**) The correlation plot showed the protein-coding gene expression levels between the ss&rd_004 and IVT_neg_004 samples. Spearman’s rank correlation coefficients were calculated using CPM normalization. Undetected genes were excluded from the analyses. Each point represents one gene, color-coded by the density at the plot position. Gene length is indicated by the point size. (**F**) Observed accuracy distribution including mapping accuracy, insertion, deletion, and substitution of ss&rd_004 and IVT_neg_004. (**G**) Mapping accuracy of site level on different RNA types.

All mapped bases were classified based on their annotation features (**Figure 1D**). In ss&rd_004, 52.3% of the total annotated bases fell within CDS, covering 3,872 genes with at least five reads each. In IVT_neg_004, 38.9% of the annotated bases were CDS regions, encompassing 3,150 genes (≥5 reads). Combined with the enhancement in sequencing yield, the number of CDS bases in both samples reached approximately 1800Mb (**Figure 1D**).

The ss&rd_004 sample and IVT_neg_004 datasets showed a high correlation in gene expression, with a Spearman correlation of 0.95 (**Figure 1E**). We observed an increased ratio of ncRNA in IVT compared to WT in both RNA004 and RNA002 datasets. This increase was likely attributable to the ncRNA *ssrA*. Previous research (16) has demonstrated that *ssrA* was highly expressed in *E. coli*, which can lead to its preferential amplification during the IVT process.

Overall, the new RNA004 kit and flowcell enabled the generation of high-yield and high-quality RNA sequencing data for both native and unmodified samples, greatly facilitating downstream transcriptome and epi-transcriptome analyses.

### The estimated and observed features of IVT surpassed those of the WT sample

Previous research has demonstrated that modifications can affect ONT raw current signals, leading to mapping errors such as insertions, deletions, and substitutions (34, 35, 46). Consequently, the unmodified control, like an IVT sample, is expected to outperform the WT sample in observed accuracy. Consistent with this, our sequencing data revealed a median accuracy of 97.5% for the WT and 98.4% for the IVT (**Figure 1F**).

A previous study showed that the accuracy varied by basecalling models version for both WT and IVT under the RNA002 kit (17). Here, we evaluated several features of mapped reads from WT and IVT across two recent basecalling models: rna004_130bps_sup@v3.0.1 and rna004_130bps_sup@v5.0.0 (**Figure S1A**). Dorado v5.0.0 model brought a significant improvement in mapping accuracy by at least 2.5% compared to v3.0.1 model. And v5.0.0 model increased the estimated Q score by approximately 7 for both WT and IVT samples, while maintaining consistent read length. Furthermore, in both versions of the basecalling models, the IVT samples demonstrated higher accuracy than the WT.

We next compared the site-level mapping accuracy among different RNA types in RNA004 dataset using Dorado v5.0.0 model. We found that the transfer RNA (tRNA) exhibited the largest accuracy gap (1.6%) between WT and IVT samples (**Figure 1G**). The tRNAs are known to harbor a high density of modifications. We believed these modifications account for the lower accuracy of tRNA in the WT sample compared to the other types.

Although Q scores were commonly used as filtering criteria in DNA sequencing, their applications in RNA studies remain relatively uncommon. For our RNA004 data, a Q score threshold of 7 was proved effective for pre-filtering unmapped reads in the v5.0.0 model (**Figure S1B**). However, this threshold may differ for other models: for example, a Q score of 5 worked better under the v3.0.1 model. To investigate the underlying significance of Q scores, we conducted a correlation analysis between the estimated Q score and the Phred-scale observed mapping accuracy. Although positive correlations were observed, the relationships were not statistically strong (**Figure S1C**). Therefore, it is recommended to analyze Q scores and observed accuracy independently in nanopore RNA studies.

In conclusion, our RNA004 data confirmed that IVT achieved higher read quality than WT, as reflected by both estimated and observed features across two basecalling model versions. The largest divergence occurred in tRNA, where the WT harbored numerous modifications. We also found that Q scores could be used to filter poor-quality RNA reads, but this approach should be tailored to each basecalling model version.

### High false positive rates were observed when using Dorado’s modified basecalling models

To evaluate the performance of Dorado modified basecalling models in *E. coli*, a series of analyses were performed based on our WT and IVT datasets (**Figure 2A**). The model of rna004_130bps_sup@v5.1.0 was compatible with m^6^A, Ψ, A-to-I, and m^5^C all-context models. Here, we ran all the all-context RNA modified basecalling with the v5.1.0 models. The Dorado modified basecaller generates a BAM file containing modification information (Modbam). Using another official tool called Modkit, the Modbam file could be converted to a BED file (Modbed) for downstream analysis. We retained sites with adequate coverage (>20) in the Modbed file and analyzed their “fraction modified”, defined as the ratio of modified nucleotides among all.

**Figure 2.**
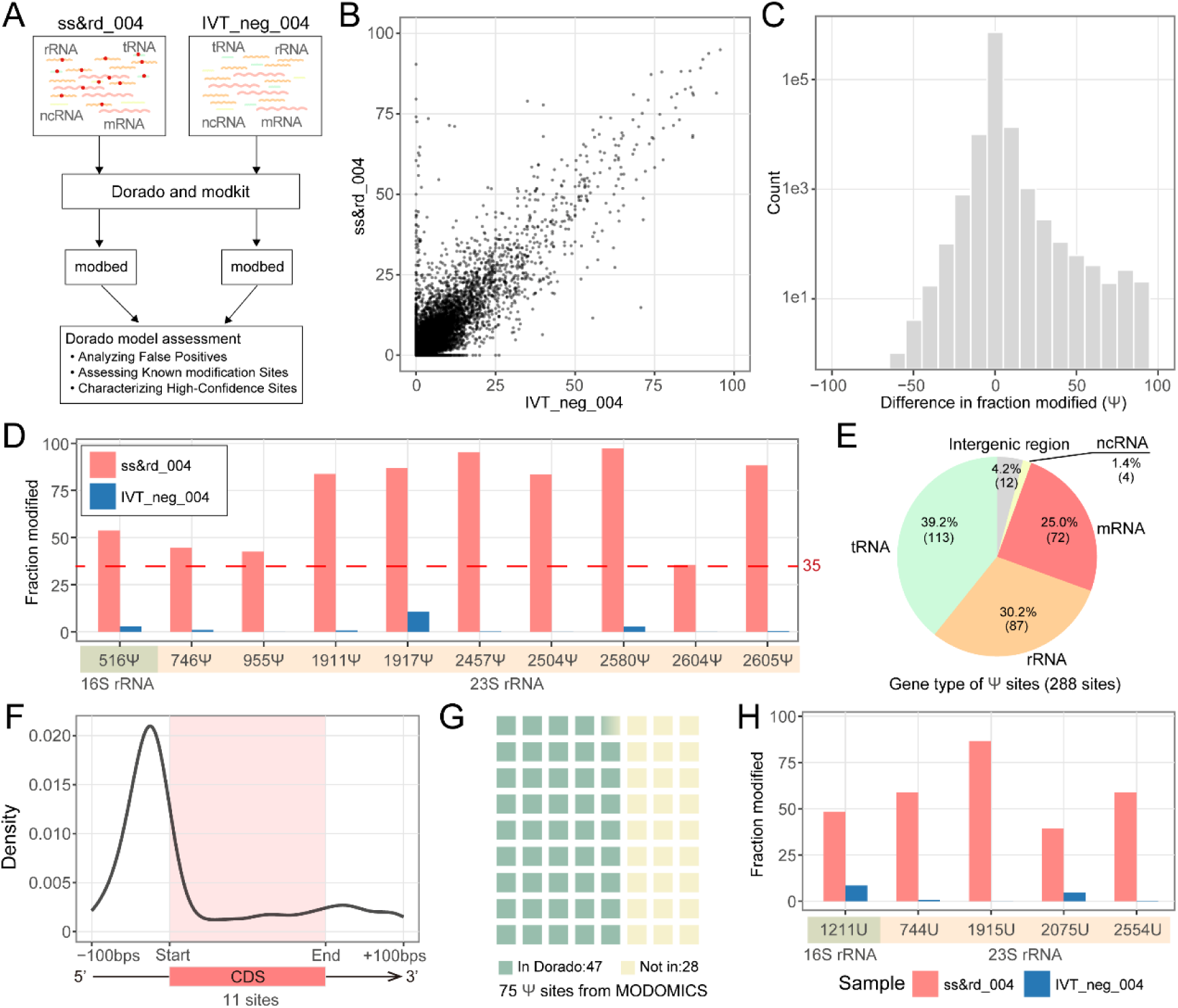
Evaluation and characterization of Dorado Ψ model. **(A)** Workflow of our analysis based on Dorado Ψ model. The raw data of ss&rd_004 and IVT_neg_004 were basecalled by Dorado v5.1.0 Ψ model. BAM files with modification information were transferred to BED files with modification information (modbed) by modkit tool. Further assessments were done for the Dorado model results. **(B)** The dotplot illustrated the “fraction modified” value of each position detected by the Dorado Ψ model from WT (ss&rd_004) and IVT (IVT_neg_004) samples. **(C)** The right-skewed distribution of difference of “fraction modified” (WT - IVT). The y-axis was treated with log10. **(D)** “fraction modified” value of 10 well-studied Ψ sites on *E. coli* rRNA. The lowest difference is around 35. **(E)** The proportions and number of high-confidence sites within the expanded region. To investigate the characteristics of high-confidence sites located in the UTR region, we expanded the annotation region by 100 base pairs at both the start and end of each gene. This adjustment enabled the CDS to encompass nearly the entire mRNA sequence. **(F)** Density distribution around the CDS region, with 11 in 72 sites located within the CDS. **(G)** The waffle chart shows the number of Ψ sites validated by Dorado in MODOMICS. 47 Ψ sites on tRNA were identified in comparison with MODIMICS. **(H)** “fraction modified” of high-confidence sites on 16S and 23S rRNA that were previously unreported.

As a negative control, the “fraction modified” values of each site in IVT_neg_004 were expected to be significantly lower than that of ss&rd_004. When analyzing the result from Ψ all-context model, 92.4% of well-covered sites in IVT_neg_004 also appeared in ss&rd_004, which can be used for further comparison. However, only 40.7% of the modified fraction from well-covered sites was lower than ss&rd_004 (**Figure 2B** and **Figure S2A**). This suggested that ss&rd_004 also may contain many false positives. When false positives are high, using a negative control can help filter them out. We calculated the difference in modified fraction by subtracting IVT from WT for each site. The average difference was close to zero, and the distribution was right-skewed (**Figure 2C** and **Figure S2B**), suggesting certain sites displayed signal patterns indicative of true positive modifications at defined thresholds.

To verify whether the differences between WT and IVT could indicate true modifications, we initially focused on 36 well-studied modified sites in *E. coli* rRNA. These sites included ten Ψ sites, two m^6^A sites, and three m^5^C sites. Utilizing the Dorado v5.1.0 model for Ψ detection, we found clear differences (at least 35%) between WT and IVT (**Figure 2D**). Setting a threshold of 35% for the difference in “fraction modified” yielded 288 high-confidence Ψ sites, with 25% on mRNA, 39.2% on tRNA and 30.2% on rRNA (**Figure 2E**). The majority of reliable sites on mRNA were enriched in the 5 prime untranslated region (5’ UTR) (**Figure 2F**). Cross-referencing Dorado’s tRNA Ψ positive sites against the 75 known tRNA sites deposited in MODOMICS database (a comprehensive public RNA modification database) identified 47 overlaps (**Figure 2G**). While this difference-based method captured known rRNA Ψ sites accurately, we also found five unreported rRNA sites (**Figure 2H**).

For the m^6^A model, the differences between WT and IVT at the two known m^6^A sites, A1618 and A2030, were more pronounced compared to those observed in the Ψ model, exceeding 65 (**Figure S2C**). Applying the same strategy, we spotted one additional unreported site on the 23S rRNA (A2448). We also identified another modified site, A1518 on 16S, which was known to be a m^6,6^A modification (47) instead of m^6^A (**Figure S2C**). Among annotated regions, most high-confidence m^6^A sites (152 in total) were in mRNA (**Figure S2D**). They clustered near the 5′ UTR (**Figure S2E**), and motif analysis with MEME revealed a strong signal in the polyadenylation A (polyA) region (**Figure S2F**).

For the m^5^C and A-to-I models, a similar distribution pattern between WT and IVT “fraction modified” value difference was observed (**Figure S2G** and **S2H**). For the three known m^5^C sites on rRNAs, the difference at C1962 on 23S rRNA is large (>75), but the differences at C967 and C1407 on 16S rRNA were minimal (<15) (**Figure S2I**). We couldn’t simply use one cutoff to differentiate the true positive site using the current m^5^C model. For the A-to-I, no known sites or ground truth exist in *E. coli* so we couldn’t determine a good cutoff. As a result, no further downstream analyses were done using m^5^C and A-to-I Dorado models. Instead, we provided the sites whose differences between WT and IVT “fraction modified” values exceeded 50 from the two models in Table S2.

In conclusion, the Dorado modified basecalling model exhibited a high false-positive rate in *E. coli*. An IVT sample was necessary to filter out false positives. Yet, certain known Ψ and m⁶A sites showed strong WT to IVT differences. This observation prompted us to leverage fraction-modified differences for high-coverage sites, thereby boosting our confidence in identifying true modification sites. Utilizing “fraction modified” differences between WT and IVT for filtering high false-positive sites could enable more detailed follow-up studies. The need to include IVT samples in Dorado’s single-mode workflow has limited its direct applications for bacterial RNA modification detection. This constraint suggested that a comparative approach—leveraging WT–IVT signal differences—could offer a more effective solution.

### nanoSundial, a new modification detection tool based on RNA004 was designed for prokaryotes

Despite the value of Dorado’s modified basecalling, our analyses showed that it can produce a high false-positive rate in *E. coli*. Including an IVT (unmodified) sample helped exclude many false positives by comparing “fraction modified” between WT and IVT, and indeed confirmed strong WT-to-IVT differences for certain known Ψ and m⁶A sites. However, even after applying this filtering step, Dorado still detected some unreported sites, raising the possibility of algorithmic artifacts or undescribed modification types. Moreover, there were more than 160 known RNA modifications (1–6), yet the Dorado v5.1.0 model only supports four (Ψ, m⁶A, m⁵C, and A-to-I). These observations underscored the need for a comparative strategy that exploits WT–IVT signal differences and can potentially identify a wide spectrum of modifications.

To address the limitations, we developed *nanoSundial*, a new comparative modification detection tool specifically tailored to prokaryotes using RNA004 data. *nanoSundial* took advantage of the improved signal-mapping accuracy in RNA004 (often referred to as resquiggle or eventalign). It then applied a statistical framework that directly compared the raw signal profiles of WT and IVT samples, rather than relying on single-mode classification from a limited subset of known modifications (**Figure 3A**; see Methods). In doing so, *nanoSundial* aimed to provide a more comprehensive and reliable approach to detecting diverse RNA modifications in *E. coli*.

**Figure 3.**
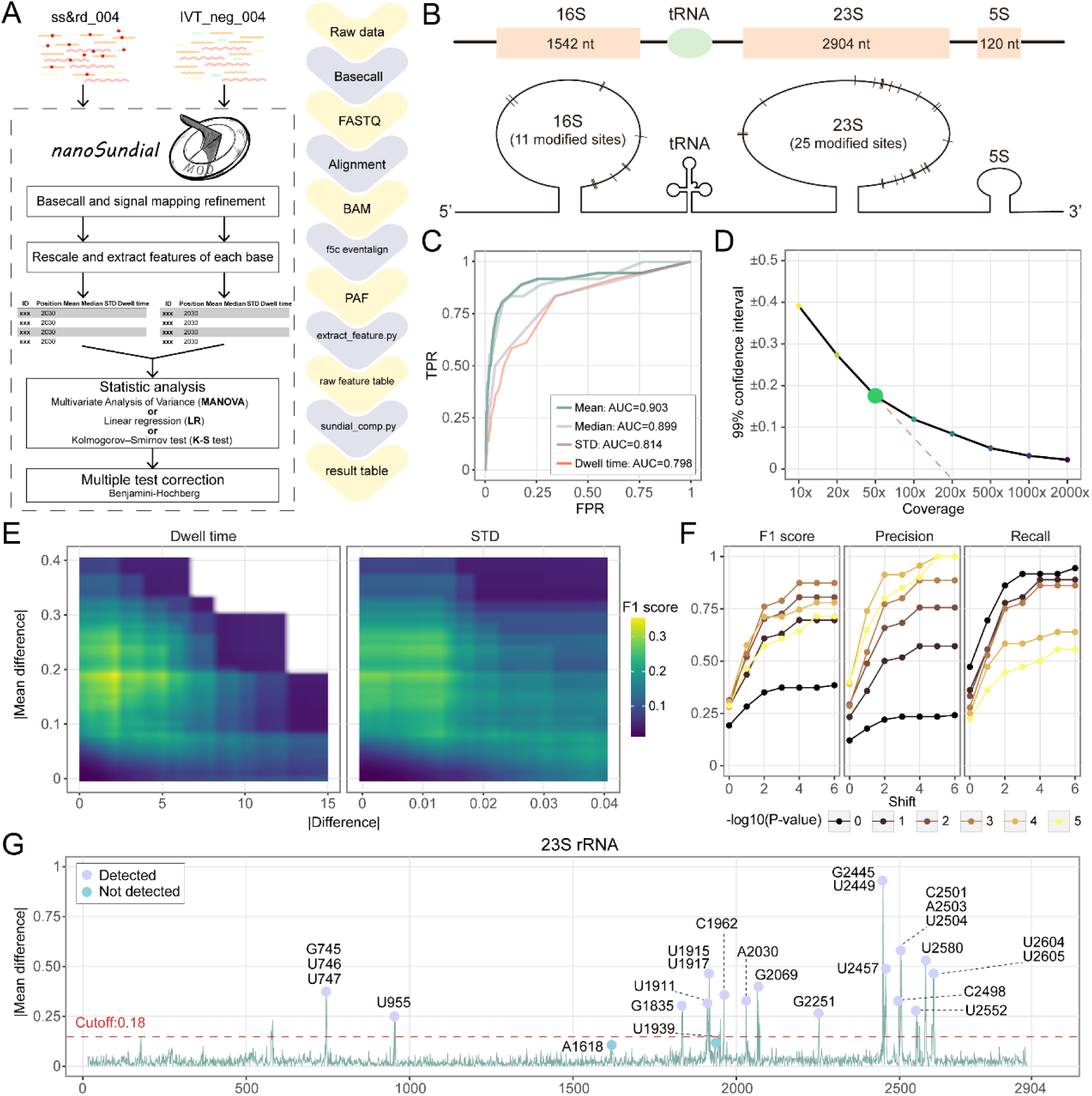
Workflow and optimization of the nanoSundial. **(A)** Raw data WT (ss&rd_004) and IVT (IVT_neg_004) were basecalled using Dorado, filtered with Samtools, and then underwent signal mapping refinement via f5c eventalign. NanoSundial then rescales the data and extracts features including mean, median, dwell time, and STD for each site. Finally, statistical analyses, including MANOVA, or LR, or the K-S test, were performed, followed by multiple testing corrections using the BH method. The right side highlights all involved programs in purple boxes and files in yellow boxes. **(B)** The diagram shows the distribution of 36 known modification sites on the E. coli rRNA, representing the ground truth. **(C)** Receiver Operating Characteristic (ROC) curves for the four features were generated based on the absolute differences between the two conditions. **(D)** The 99% confidence interval of the absolute difference in mean when comparing two independent negative controls at different coverage levels. This value shows a sharp change in slope at a coverage of 50x. **(E)** F1 score distributions when applying the absolute mean difference combined with the absolute differences in dwell time or standard deviation as co-cutoffs. The highest F1 score is achieved when the absolute difference of dwell time is around 1 and the mean is around 0.18. **(F)** Dot and line plots illustrate the variation in F1 score, precision, and recall (on the y-axis) as different cutoff values of adjusted *p*-value are applied, with the x-axis representing the shift length at a 50x coverage. The highest value is observed when -log10 adjusted *p*-value reaches 3, and the results stabilize when the shift is set to 4. **(G)** Plot shows the absolute mean difference on the 23S rRNA. The purple points represent the sites within a 4-base shift that can be detected by nanoSundial, while the blue points indicate those that were not detected.

We used 36 well-characterized modification sites on *E. coli* rRNA (**Figure 3B**) as true positive sites. These included 11 sites from the 16S rRNA and 25 sites from the 23S rRNA. We generated receiver operating characteristic (ROC) curves to compare the classification performance using absolute differences in mean, dwell time, and standard deviation (STD) between ss&rd_004 and IVT_neg_004. Among these features, the absolute mean difference achieved the highest area under the curve (AUC) score of 90.3% (**Figure 3C**), while all four features surpassed 79.8%.

Fine-tuning the cutoff across various coverage levels has consistently posed a challenge for comparative tools (48). To minimize false positives, we evaluated our method on two non-overlapping subsets of IVT_neg_004 (negative controls) with coverage levels from 10x to 2000x (**Figure 3D**). Coverage had a major impact. As coverage increased, the 99% confidence interval narrowed. A notable change in slope occurred at 50x coverage, after which the slope became more gradual. And the 99% confidence interval at higher coverage levels was close to zero. For RNA004 data, we recommend a minimum coverage of 50x, similar to the Tombo_level approach (49) used for RNA002.

Though the mean difference between IVT and WT is the key factor for RNA modification detection, analyzing the differences in dwell time and STD on the 23S rRNA demonstrated their possible contribution (**Figure S3A, S3B**). We would like to know if we could include one additional factor to achieve a better detection result. So we evaluated the F1 score across various cutoff values using the mean together with Dwell time or STD (**Figure 3E**). Combining STD with mean did not enhance the F1 score, but including dwell time did. The highest F1 score (0.34) was achieved with an absolute mean difference cutoff of 0.18 and an absolute dwell time difference cutoff of 1. These cutoffs offered an optimal balance between precision and recall. Consequently, we recommend a cutoff of 0.18 for the absolute mean difference and a cutoff of 1 for the absolute dwell time difference.

Determining an appropriate cutoff for the *p*-value posed another challenge (48). We observed that, above 200x coverage, the adjusted *p-*value from MANOVA lost effectiveness (**Figure S3C**). However, at low coverage (50x), the adjusted *p*-value remained highly informative, with the best performance at the -log10 (adjusted *p*-value) = 3 (**Figure 4F**). Therefore, we recommend a *p-*value of 1e-3 as a universal cutoff across coverage levels when using MANOVA.

**Figure 4.**
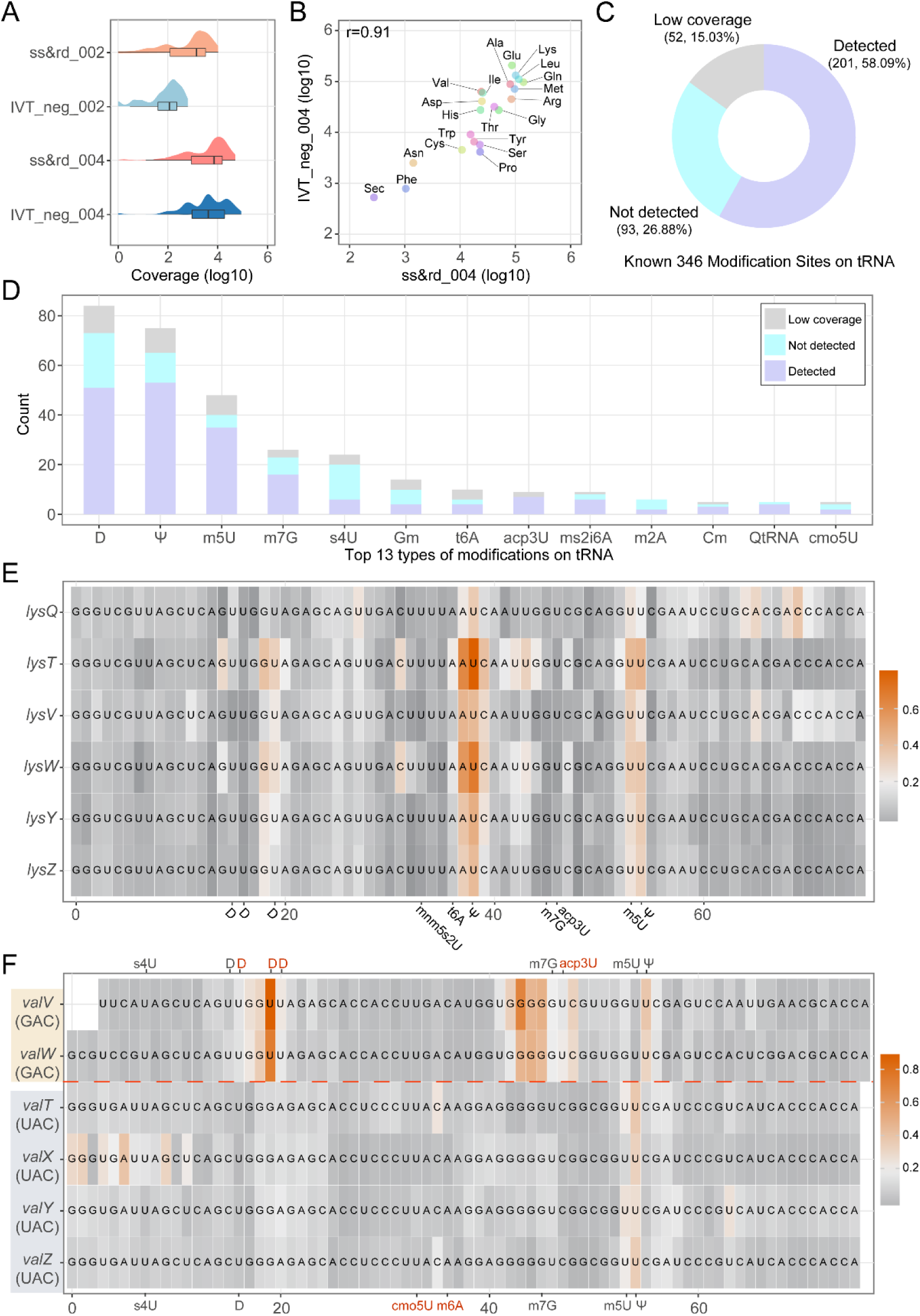
Characterization and validation of nanoSundial in tRNA applications. **(A)** Coverage (log10 scale) on tRNA for the RNA004 data compared with the RNA002 data. **(B)** Correlation between WT (ss&rd_004) and IVT (IVT_neg_004) on tRNA expression levels. **(C)** The performance of nanoSundial using MODOMICS as the ground truth, detecting 201 out of 346 sites (58%), with 52 positions (15%) having coverage below 50x. **(D)** Top 13 types of modifications on tRNA. **(E-F)** Heat maps illustrate the absolute mean difference for the paralogs of tRNA lysine (lys) (E) and valine (val) (F). The colors at each position represent the magnitude of these absolute differences. For valine, the paralogs can be categorized into two classes based on their binding codons (GAC and UAC), which also exhibit different modifications. These different modifications are highlighted in the red text.

Unlike tools specifically designed for m^6^A modification detection, *de novo* modification detection methods lack prior knowledge of the motifs associated with each modification type. Additionally, modifications can alter the current signals in neighboring nucleotides (32, 34). Thus, a shift strategy is necessary to identify potential modification sites, similar to earlier pipelines (17, 34). Using our recommended cutoffs, performance plateaued at the shift of 4 bases (**Figure 3F**), indicating no further gains beyond that point.

After all optimizations, this approach successfully identified 34 out of 36 known rRNA sites (**Figure 3G**, **S4A**, and **S4B**). Aside from the detection of known positive rRNA sites on 16S and 23S rRNA, our analysis on 5S rRNA, which held no modification, found no positive hits, signifying reliable performance for true negatives. The two missing positive sites were A1618 and U1939 on 23S rRNA. A1618 may not be fully modified, and m⁵U might not produce a signal change comparable to other modifications.

In summary, we developed nanoSundial—a modification detection tool optimized for bacterial data from the RNA004 kit. To optimize our approach, we employed 36 known modification sites on rRNA as ground truth. We assessed various features, including mean, median, standard deviation, and dwell time. By optimizing coverage thresholds, cutoff values, and shifting, nanoSundial demonstrated strong performance in identifying modifications on rRNA.

### The performance of nanoSundial was validated in tRNA modification sites

Compared to RNA002, RNA004 data had a marked increase in sequencing yield per flowcell, accompanied by substantial improvements in site-level coverage (**Figure 4A**). The median coverage for tRNAs now exceeded 2,000 in RNA004. Meanwhile, tRNA expression levels in ss&rd_004 and IVT_neg_004 were highly similar (**Figure 4B**), with a correlation coefficient of 0.91. The enhancements in both sequencing depth and the expression level concordance on RNA004 provided a solid foundation for comprehensive tRNA modification analysis.

We drew upon 346 modification sites from MODOMICS (6) as the ground truth. By employing a 4-base shift strategy, nanoSundial successfully detected 201 of these sites (**Figure 4C**). Focusing exclusively on tRNAs with sufficient sequencing depth in our dataset, we achieved a 68.3% detection rate for known tRNA modifications, belonging to 34 types of tRNA modifications. We also calculated site detection ratios for the 13 most common tRNA modification types (**Figure 4D**). Of these 13 types, nanoSundial detected over half the sites in 9 types.

nanoSundial exhibited stable performances across tRNA paralogs. As an example, we examined mean WT to IVT differences in lysine (lys) tRNAs (**Figure 4E**), which include six paralogs. The distribution of mean differences was remarkably similar across all paralogs. In each case, Ψ sites exhibited the greatest mean differences compared to other modifications (**Figure 4E**). The example of one lys paralog (*lysW*) further indicated the dramatic change of current signals between WT and IVT around modified sites (**Figure S5A**). Additionally, we provided mean differences for another tRNA with multiple paralogs, aspartic acid (asp) (**Figure S5B** and **S5C**). All these showcases demonstrated nanoSundial’s capability of detecting modifications on tRNAs.

nanoSundial effectively distinguished the modification profiles among tRNA paralogs that bind to different codons. Identical tRNA types may exhibit distinct modifications depending on their codon binding. We further analyze multi-codon tRNA, such as tRNA^Val^, which binds GAC or UAC (**Figure 4F**). Specifically, *valV* and *valW*, which pair with the GAC codon, possess three additional 5,6-dihydrouridine-5’-monophosphate (D) modifications and one additional 3-(3-amino-3-carboxypropyl) uridine (acp3U) modification compared to their paralogs that binding the UAC codon. As a result, regions adjacent to these additional modification sites displayed higher absolute mean differences in *valV* and *valW* (**Figure 4F**). These findings pointed out nanoSundial’s ability to differentiate the modification profiles of tRNA paralogs with different binding codons.

The improved sequencing data provided by RNA004 has facilitated the analysis of tRNA. nanoSundial not only detected a greater number of modifications but also identified modifications across different codons within tRNAs.

### Consistent modification detection performance of nanoSundial across RNA types

Modified bases can influence the nanopore raw current signals for neighboring regions (26). In our analysis, nanoSundial output all sites satisfied the filter of current differences, including both the modified sites and neighboring sites. To refine these outputs, we employed a merging strategy to consolidate adjacent sites into positive regions (**Figure 5A**; see Methods).

**Figure 5.**
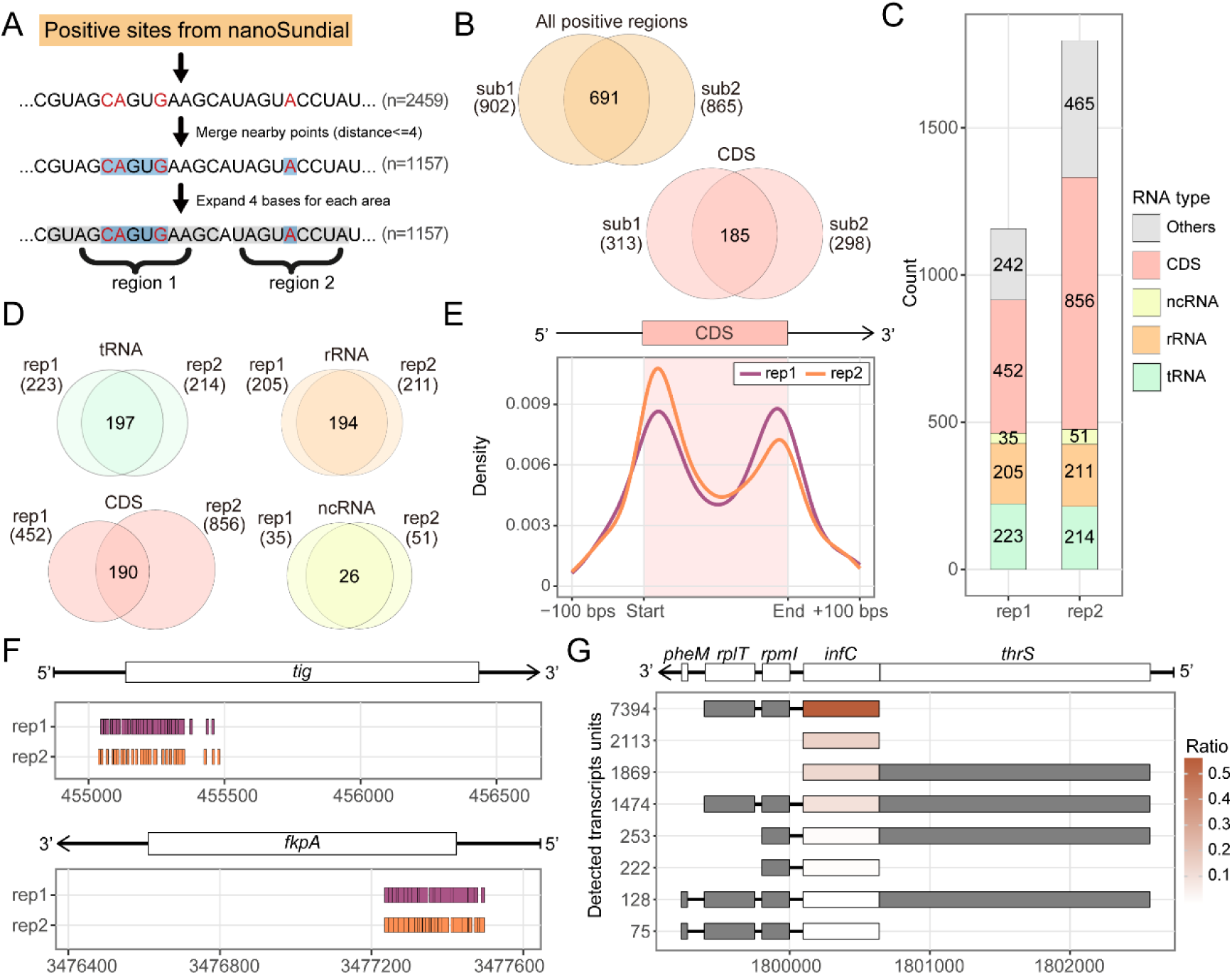
Reproducibility testing and high-confident positive region exploration of nanoSundial. **(A)** The merging strategy will connect all nearby positive sites within 4 bps, then expand by an additional 4 bps, ultimately identifying and outputting these positive regions. **(B)** Venn plots show the intersection of overall and CDS between the results from two non-overlapping subsamples (technical replicates). **(C)** The barplots demonstrate the number of positive regions within the annotation feature of two biological replicates result. Numbers embedded in the bars indicate the amount of positive regions in each category, including tRNA, rRNA, ncRNA, CDS, and others. Others covered the UTR region and intergenic region. **(D)** The intersection of positive region on each category of gene type. **(E)** Density distribution around the CDS region of two biological replicates result. **(F)** Showcase of the clustering of positive regions on the genes *tig* and *fkpA*. Positive regions were enriched in the start of mRNAs. **(G)** The composition and proportion of the Transcription Units (TU) of the operon containing *infC*. The color indicates the proportion of each TU relative to the total reads across gene *infC*. The TU with the highest expression has *infC* as its first gene.

Reproducibility remains a major challenge in modification detection. Previous studies suggested multiple replicates (32). To evaluate the reproducibility of nanoSundial, we conducted both technical and biological replicate analyses. For technical replicate analysis, WT and IVT were divided into two non-overlapped halves. Each half of the WT sample was then compared with one half of IVT, resulting in two sets of results from nanoSundial, referred to as sub1 and sub2. (**Figure S6A**). By applying the coverage cutoff of at least 50, approximately 4% of the sites in one subset (sub1 or sub2) were absent in the other subset and were therefore excluded (**Figure S6B**). Sub1 returned 902 positive regions, while sub2 returned 865. Overall, their intersection consisted of 691 regions, accounting for approximately 76% of sub1 and 78% of sub2. The intersection rate for mRNA was the lowest, which is less than two-thirds (**Figure 5B**). In contrast, the intersection rates for tRNA, rRNA, and ncRNA exceeded 95% (**Figure S6C**). This discrepancy suggested that many modifications on mRNA may be only partially modified, resulting in differences between WT and IVT mRNA that were insufficient to be reliably detected at certain modification points. This issue became more pronounced when sequencing coverage was reduced through subsampling, making the differences even less significant.

We then explored biological reproducibility by sequencing an additional biological replicate of ss&rd_004 (WT). The size selection step before rRNA depletion and polyadenylation may result in the different proportions of small RNA, rRNA and mRNA between batches. Nonetheless, the replicate showed high yield and quality, with sequencing yield exceeding 6.2 Gbps and median of Q score > 17.93 (**Table S1**). The correlation of gene expression between the two WT replicates is 0.98 (**Figure S6D**).

Both WT biological replicates were compared to the IVT_neg_004 sample using nanoSundial. After applying the merging strategy, rep1 yielded 1,157 positive regions from the initial 2,459 positive sites, and rep2 yielded 1,797 positive regions from 3,512 sites (**Figure 5C**). Most positive regions were in the coding sequence (CDS), with 452 for rep1 and 856 for rep2. Overall, rep1 and rep2 shared 741 overlapping regions. The intersection rates were especially high for tRNA and rRNA. For example, 194 of ∼200 rRNA regions overlapped (**Figure 5D**). Despite 452 regions in rep1’s CDS and 856 in rep2’s, only 190 overlapped, indicating a lower intersection rate in mRNA regions.

In summary, we incorporated a merging strategy and demonstrated through both technical replicate and biological replicates that nanoSundial had good reproducibility, especially on tRNA and rRNA regions. Moreover, in two biological replicates, we detected 190 intersecting positive regions within the CDS. These regions were of higher confidence compared to other positive regions in the CDS, making them prime candidates for downstream analysis.

### mRNA modifications were enriched at the start and end of CDS

By analyzing the density of all positive regions located on mRNA, we found that modifications were primarily concentrated within the gene body. The enriched peaks of modification regions were close to the CDS start and end (**Figure 5E**). This distribution was consistent in the two biological replicates. While observing the distribution characteristics of positive regions, we noticed a phenomenon that many nearby regions were enriched in one mRNA.

We further analyzed the location of the 190 high-confident positive regions in the CDS area. The 190 overlapping regions involve 71 genes, with operon information from regluonDB described in Table S3. We defined the 71 genes as stably modified genes. Although 55 out of 71 genes have only one or two positive regions, some mRNAs, like *tig*, *fkpA*, *cutC*, *sbmA* and *OmpC*, exhibited a clustering of adjacent regions at around the 5’ or 3’ ends (**Figure 5F**).

Among the 71 genes, we found that 25 genes are part of operons containing three or more genes, and 22 in 25 genes are predominantly located at the beginning or end of the operons (see **Table S3**). Notably, some operons that start or end with a stably modified gene, like *xerD*, *lptD*, *thrS*, *rplQ*, and *rpsQ*, contain more than five genes. In particular, the *rpsQ* operon consists of 11 genes. Within the 71 stably modified genes, only 3 genes, including *infC*, *secD*, and *yggX*, were not located at the start or end of the containing operon. Further analysis of their TU compositions revealed that the three genes were positioned at the start or end of the most abundant TUs (**Figure 5G**, **Figure S6E**, and **S6F**) in the operon.

Each of the highly expressed TUs comprised more than 85% of the total expression for each operon.

We investigated the biological functions of operons (with more than three genes) that contained stably modified genes. Among these operons, nine had a stably modified gene located at the start of the operons, encompassing a total of 42 genes, while eleven had a stably modified gene at the end of the operons, accounting for 50 genes. Both sets were enriched in the “translation” Gene Ontology (GO) pathway (**Figure S7**). Notably, the operons with stably modified genes at the ends were also associated with multiple “ribosome”-related functions. These findings suggested that operons with stably modified genes at the start play key biological roles in translation, with those with stably modified genes at the end may have broader implications for ribosome biology.

In summary, our biological replicates confirm that positive regions on mRNA tend to cluster near the CDS boundaries. Moreover, the 71 stably modified genes typically lie near the start or end of their TUs, suggesting a potential functional significance for modification sites in these regions.

## Discussion

In recent years, substantial progress has been made in understanding the roles and functions of RNA modifications. Modifications in eukaryotes, such as m^6^A and m^5^C, are well characterized, but bacterial mRNA modifications remain less understood (50). The latest nanopore RNA sequencing kit (RNA004) had great potential to tackle this gap, as it provides higher yield, lower signal to noise ratio, and higher-quality reads compared to the previous RNA002. Several computational methods, including single-mode and comparative methods, were originally designed for RNA002. To date, only the Dorado modified basecalling model has been developed for the RNA004, it represents a single-model method that does not require negative controls and can analyze modification stoichiometry at the site level.

In this case, we applied the latest RNA004 technology to detect RNA modifications in *E. coli*. We conducted an improved experimental pipeline and prepared sufficient yields of both native (wild type, WT) and unmodified (*in vitro*-transcribed, IVT) RNA samples. Combined with the negative control (IVT), we evaluated the modification detection models provided by Dorado (Ψ, m^6^A, m^5^C, and A-to-I) on our *E. coli* data and found a large number of false positives. As a result, we concluded that the Dorado’s direct-RNA single-model modification basecalling models couldn’t be directly applied to prototypes without using a negative control to filter away the false positives. By observing the “fraction modified” values of known modified sites on rRNA, we proposed a simple pipeline to identify the high-confidence sites using the difference in “fraction modified” between WT and negative control. After optimization, the Dorado pipeline could successfully identify all known Ψ and m^6^A sites. However, even with the improvement with IVT, it detected additional Ψ and m^6^A sites on rRNA that have not been previously reported, which are likely to be false positives. What’s more, there were more than 160 types of RNA modifications, but the Dorado v5.1.0 model can only detect 4 types. These limitations pointed to the necessity of launching a new modification detection tool to detect bacteria.

Apart from the limitations of Dorado, previous RNA modification detection methods based on RNA002 were not compatible with RNA004 data. Furthermore, the modifications in prokaryotes differ from those in eukaryotes and do not follow the RRACH motif. Hence, we aimed to develop a comparative method. Comparative methods required a low modification or no modification sample as a negative control. Existing RNA002-based comparative methods rely on either current features or alignment features (37). Previous research showed that alignment features, etc. mismatches, highly depended on the basecaller version (17). Additionally, modifications like m^5^C and m^4^C often produce subtle or negligible current shifts, making them difficult to classify as basecalling errors (16, 34). As RNA004 improved mapping accuracy, the mismatch differences decreased between WT and IVT, making alignment features even less reliable. For example, differences at A1618 and A2030 (**Figure S8A, S8B**) are smaller in RNA004 than in RNA002 (28). Current features, on the other hand, remain largely unaffected by improved basecalling accuracy. So, we think that the comparative method with current features could better detect modification compared to methods using alignment features in bacteria. Therefore, we designed nanoSundial, which is a comparative modification detection tool for prokaryotes that specifically targets current features in RNA004 data.

The comparative strategy of nanoSundial is training-free and can potentially detect any RNA modification, so long as a negative-control sample is available. Moreover, it can identify a wider range of modifications than single-mode tools. Despite its advantages, nanoSundial also has some drawbacks. First, it is challenging to accurately pinpoint the exact base of a modification and instead locate the broader region (±4 bases). Despite the challenge in base shifting, nanoSundial detected more tRNA modification sites (201 in 346) with narrower shifts compared to a previous RNA002 kit pipeline, which identified bacterial tRNA with shifts of ± 10 bases (198 in 346) (16). This comparison indicated the improvement of RNA004 enhanced the modification base assignment. Second, compared to the single mode, it is hard to calculate the stoichiometry of modifications at the site level, which complicates the detection of partial modifications. Third, our ground truth for cutoff selection includes 36 known rRNA-modified sites, most of which are highly modified. It is noteworthy that there are over 160 different types of modifications, some of which may induce only subtle current changes between WT and IVT. The linear cutoff applied in nanoSundial may be too harsh for specific types of modifications. We could further improve nanoSundial by applying kmer specific cutoff after obtaining more ground truth.

After optimizing on rRNA and validating on tRNA, we employed a strategy to merge adjacent sites and test the reproducibility of nanoSundial. Although the reproducibility rates for tRNA, rRNA, and ncRNA exceeded 95%, the rate for mRNA was only 61%. Additionally, the replicate results showed a high intersection on rRNA and tRNA, but not on mRNA. This indicated that the modifications on mRNA are partial and unstable, consistent with observations in rRNA, tRNA, and snRNA sites from humans and yeast (10).

Still, we detected 190 positive regions from the replicate intersections, spanning 71 stably modified genes on CDS, which represent high-confidant mRNA modifications. On the gene level, 68 of 71 stably modified genes were located at the start or end of the operon. For 3 genes that did not satisfy this regulation, we found these genes were located at the beginning or end of the most highly expressed transcription unit (TU). This suggested that mRNA modifications were closely related to operon structure, which deserved further verification and study. Besides, among these 3 genes, we observed that the *napF* operon contains 15 genes. However, in the TU research, only three genes—*napA*, *napD*, and *napF*—were identified. Notably, *napF* also conforms to the positional pattern observed at the end of the operon. This discrepancy may suggest that the operon annotations in RegulonDB could be further refined, highlighting the potential application of ONT DRS in transcriptome analysis (**Figure S9**).

In addition, we assessed nanoSundial’s consistency with Dorado in detecting other modifications. Comparing high-confidence Ψ sites from Dorado with nanoSundial revealed a substantial overlap (177 out of 288 sites), whereas the m⁶A all-context model performed poorly, intersecting at only 38 out of 234 sites (**Figure S10A**). This finding indicated a high level of consistency between nanoSundial and Dorado Ψ model, mainly for rRNA and tRNA (**Figure S10B**). By contrast, the lower overlap for m⁶A likely reflects limited accuracy in Dorado’s all-context m⁶A model, and the presence of partial modifications in m⁶A —such as A1618—which nanoSundial struggled to detect. The high Ψ overlap and low m^6^A overlap between Dorado and nanoSundial suggest that Dorado may produce more false positives in bacterial m^6^A detection.

In conclusion, the enhanced quality and yield of RNA004 data have substantially supported our analysis of bacterial RNA. We first utilized the RNA004 data to evaluate the performance of the Dorado modified basecalling models and found that Dorado produced a high number of false positives when detecting bacterial RNA modifications. Although we implemented a method to effectively reduce most of these false positives, some unreported modified sites remained. To address the limitations of the Dorado model and to establish a tool capable of precisely detecting RNA modifications in bacteria, we developed nanoSundial, a new comparative method for prokaryotic RNA004 data based on current features. Following our optimizations and evaluation, nanoSundial not only demonstrated strong performances in detecting RNA modification in bacterial tRNA and rRNA, but also revealed a preferential enrichment of modifications in bacterial mRNA. This work opens new avenues for investigating bacterial mRNA modifications at the transcriptome scale.

## Methods

### Bacterial RNA preparation

*E. coli* strain K-12 was purchased from the American Type Culture Collection, VA, USA. The preparation steps for native RNA from *E. coli* K12 were performed as described in a previously published study (17). Following the extraction of total RNA from bacterial cultures at an OD600 of 0.4–0.6 using Trizol, we purified 20 micrograms (ug) RNA with a concentration of 200 nanograms per microliter (ng/μl), using 0.8 volume of SPRIselect Beads (Beckman Coulter, IN, USA) to remove small RNA fragments (size select). Subsequently, ribosomal RNA in size selected RNA was depleted with RiboMinus™ Transcriptome Isolation Kit, bacteria (Invitrogen). The amount of size selected RNA for ribosome-depleted was 15 ug per reaction. Poly(A) tailing was conducted to the ribosome-depleted RNA using E. coli Poly(A) Polymerase (New England Biolabs, MA, USA). The native RNA of preparation was completed.

For the generation of *in vitro* transcribed (IVT) RNA, the poly(A)-tailed RNA underwent reverse transcription to synthesize double-stranded cDNA, which was then used for *in vitro* transcription. Primers used in reverse transcription referred to previously published protocols (17, 51). The following oligonucleotides were purchased from the BGI Genomics, SZ, PRC: template switching oligo (TSO) T7 primer, 5′-ACTCTAATACGACTCACTATAGGGAGAGGGCrGrGrG-3′, where r indicates ribonucleotide bases; T7 extension primer, 5′-GCTCTAATACGACTCACTATAGG-3′. For the reverse transcription conditions, we referred to a previously published literature (52). In total, 300 ng RNA in 6 μl nuclease-free water was first annealed with 1 μl oligo(dT)_23_VN primer (10 μM, New England Biolabs) and 1 μl dNTP (10 mM, New England Biolabs) for 3 min at 72°C, 10 min at 4°C, 1 min at 25°C with the lid temperature set at ≥85°C and then held at 4°C. The reverse transcription (RT) mix was assembled containing 3 μl nuclease-free water, 4.4 μl 5xRT Buffer, 1 μl RNaseOut (Invitrogen), 1 μl TSO T7 primer (75 μM), and 1 μl Maxima H Minus Reverse Transcriptase (Thermo Scientific). After adding the RT mix to the annealed RNA, the reaction mixture was incubated following the SSIV RT protocol (52). The RNA template was subsequently hydrolyzed by the Thermostable RNase H (New England Biolabs) according to the manufacturer’s instructions. The template-switching cDNA product was purified using the VAHTS RNA Clean Beads. The second-strand cDNA synthesis reaction mixture was assembled on ice consisting of 20 μl template-switching cDNA, 25 μl Q5 Hot Start High Fidelity Master Mix (New England Biolabs), 3.75 μl T7 extension primer (50 μM), and 1.25 μl nuclease-free water. Following the initial denaturation at 95°C for 1 min and 57°C for 30 sec for annealing, the reaction mixture was incubated at 65°C for 10 min. The resulting double-strand DNA (dsDNA) was purified using 1 volume of AMPure XP beads (Beckman Coulter). The *in vitro* transcription step was performed using the MEGAscript Kit (Ambion, MA, USA) at 37°C for 4 h. IVT RNA prepared in this article refers specifically to modification-free RNA, so the reaction mixture was composed of 2 μl each of NTPs, 2 μl Reaction Buffer, 120 ng ds-cDNA template in 8 μl nuclease-free water, and 2 μl Enzyme Mix.

### Nanopore direct RNA sequencing and data processing

The libraries of native RNA and IVT RNA were constructed following the manufacturer’s instructions using the SQK-RNA004 Kit (ONT, Oxford, UK). The optional RT step was performed. Nanopore direct RNA sequencing (DRS) was conducted on the MinION platform using the FLO-MIN004RA Flow Cell (ONT). The resulting POD5 files were basecalled using the Dorado workflow (v0.8.0) with the model of rna004_130bps_sup@v3.0.1 and rna004_130bps_sup@v5.0.0. The basecalling results were stored as FASTQ files and were statistically analyzed with SeqKit v2.3.0 (53). Raw read features, including read length and Q score, were extracted using Giraffe v0.2.3 (54). Subsequently, reads were aligned to the *E. coli* genome (GenBank Accession Number: NC_000913.3) using minimap2 v2.17 with parameter settings “-ax map-ont” (55). Mapping results (SAM files) were converted into BAM files and sorted and indexed using SAMtools v1.13 (56). Read alignment information was extracted from the sorted BAM files and converted into BED files using a custom Python script based on Pysam (https://github.com/pysam-developers/pysam). And accuracy feature in site level is used Epinano_Variants.py from Epinano v1.2 (28)

For the modification model, we applied rna004_130bps_sup@v5.1.0, covering three modifications models: m5C, inosine_m6A, Ψ. After the modified basecalling, the output modbam file will be aligned with the reference genome and then sorted by SAMtools. The new sorted bam files were converted into modbed files with modkit (https://github.com/nanoporetech/modkit).

### Gene expression analysis

Gene expression correlations were subsequently calculated based on the mapping results. For DRS data, reads that did not match the strand in annotation were filtered, followed by counting the read numbers aligned to individual genes. Genes other than Protein-coding regiones were removed from the counting results. Undetected genes were also excluded from the analysis. Afterward, counts per million (CPM) were computed for each gene in different samples using the formula, followed by pairwise calculation of Spearman’s rank correlation coefficients.

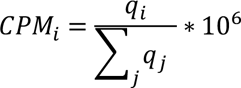

Where *q_i_* denotes reads mapped to a gene, and ∑_*j*_*q_i_* corresponds to the sum of mapped reads to individual gene. The above process was implemented using our custom script count_read_num_each_gene.py, which is now publicly available in our git and zenodo repository.

### Current feature comparison with nanoSundial

NanoSundial is a Python package dedicated to comparative analysis of DRS nanopore sequencing raw signal to de novo identify RNA modification sites. As a comparison method, nanoSundial requires two samples, a wild-type sample and a low modification or no modification as negative control sample. The analysis flow is divided into three steps (Figure 3A) after pre-processing: (1) extract current features, (2) parallel processing and statistical testing, (3) Identify the positive sites and merge the positive region.

### Pre-processing

To begin with, prepare the input data. Basecalling should first be run to obtain FASTQ files. We have chosen BLOW5 format (57), which has been proven to be smaller and delivers consistent improvements across different computer architectures. Therefore, raw data needs to be converted to BLOW5 format. After processing the two samples, an appropriate reference needs to be selected, either genome or transcriptome.

#### 1) Extract current features

The signal mapping refinement capabilities of the 004kit dataset is from f5c eventalign (58), with parameter “--pore 004kit --rna --paf --min-mapq 0”. A PAF file recording alignment information and signal index will be generated from f5c. Then, our script traverses the PAF file to extract the raw signal array from BLOW5. A thresholding normalization approach will be implemented, where the raw signal corresponding to a mapped section of each read is normalized using median shift and MAD (median absolute deviation) scale parameters.

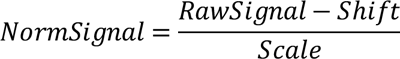

Any normalized signal values exceeding a predetermined threshold (Default: ±5) will be replaced by the threshold value. Then, for each nucleotide in each read, based on the index in the PAF file, we can obtain its window and calculate the mean, median, dwell time, and standard deviation (STD). Additionally, f5c eventalign applied a k-mer model method, there will always be some shift. Therefore, we perform the shifting operation using the shift table defined by Squigualiser (59).

Finally, since the current changes are very unstable at the beginning and end of the sequencing read, each read will by default skip the first and last ten bases (can be changed by using option “--flip”). The current characteristics corresponding to each remaining base, including mean, median, dwell time, and standard deviation (STD), will be stored on the hard drive. To reduce memory usage and speed up subsequent analysis time, a chunking strategy is used, with the default segment size being 100k bases.

#### 2) Parallel processing and statistical testing

For each position in the reference, the four features of the two samples will be compared by statistical methods. Because statistical methods can yield significantly different results when the data sizes are unequal, the “--balance” option is used to subsample the larger dataset so that the comparison numbers are equivalent.

We employ three methods to do statistical analysis, including multivariate analysis of variance (MANOVA) for all four features, as well as logistic regression and the Kolmogorov–Smirnov (K-S) test specifically for the mean to analyze and return the *p*-values. To enhance processing speed, multiprocessing has been incorporated into our script to handle each position during statistical testing. Subsequently, multiple test corrections using the Benjamini-Hochberg algorithm will be applied to adjust the *p*-values. Additionally, for each position in the reference, the four features of each sample will have their average values calculated, and then the differences between these average values will be determined. These differences, along with each position’s coverage and the *p*-values and adjusted *p*-values, will be saved in the output file.

#### 3) Identify the positive sites and merge the positive region

Through several optimizations on rRNA, appropriate cutoffs were determined as follows: an absolute value of 0.18 for the mean difference, 1 for the dwell time difference, and 3 for -log10(adjusted *p*-value). Our script will output all high confidence sites using these cutoffs and apply a merging strategy to combine adjacent points. The merging strategy consists of two steps. First, all adjacent high-confidence sites (less than 4 bases apart) are connected. Second, all points, including both those that are connected and those that are not, are extended by 4 bases to form contiguous regions.

### Reference selection and annotation strategies

When using the Dorado model, we used the genome as a reference and converted the annotation file (GFF) to a gene.bed (BED) file using a self-customized script. Then, we used Bedtools with the command “bedtools intersect -a <MODBED> -b gene.bed -wb -s”. Since the GFF file does not include UTR information and contains only the coding sequences (CDS), we will expand the annotation region of the gene by 100 bases upstream and 100 bases downstream in our study required to focus on the UTR regions.

When we optimized nanoSundial for rRNA, we used the rRNA sequences as references: 23S rRNA (GenBank Accession Number: NR_103073.1), 16S rRNA (GenBank Accession Number: NR_103073.1), and 5S rRNA (GenBank Accession Number: X00414.1). For the tRNA and other analyses, the reference genome was used (GenBank Accession Number: NC_000913.3).

### Evaluation Metrics Calculation

To assess the performance of Dorado model and optimize nanoSundial, we calculate the following evaluation metrics: TP, TN, FP, FN, Recall, Precision, F1 Score, and AUC-ROC. True Positive (**TP**): These are the instances where the model correctly predicts the positive class. True Negative (**TN**): These are the instances where the model correctly predicts the negative class. False Positive (**FP**): These are the instances where the model incorrectly predicts the positive class. This is also known as a “Type I error.” False Negative (**FN**): These are the instances where the model incorrectly predicts the negative class. This is also known as a “Type II error.”

**Recall**: Recall, also known as sensitivity, measures the proportion of actual positives that are correctly identified by the model.

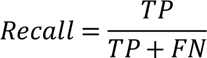

**Precision**: Precision, or positive predictive value, measures the proportion of positive predictions that are correct. It is calculated by dividing the number of true positives by the sum of true positives and false positives.

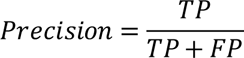

**F1 Score**: The F1 Score is the harmonic mean of Precision and Recall, providing a single metric that balances the two. It is particularly useful when there is an uneven class distribution. It is calculated by multiplying Precision and Recall by two, then dividing the product by the sum of Precision and Recall.

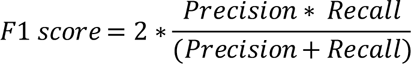

**AUC-ROC**: The Area Under the Receiver Operating Characteristic (AUC-ROC) curve measures the model’s ability to distinguish between classes. The ROC curve was constructed using two key performance indicators: the True Positive Rate (TPR) and the False Positive Rate (FPR). The TPR, also known as sensitivity, indicates the model’s ability to correctly identify positive instances. The FPR represents the proportion of negative instances that are incorrectly classified as positive.

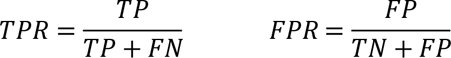

The AUC-ROC is the area under this curve, which can be calculated by integrating the ROC curve as a function of the threshold.

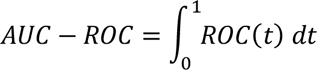

And *ROC*(*t*) is the ROC curve as a function of the threshold *t*. In our analysis, *t* represents the threshold on the absolute difference of 4 features between WT and IVT, specifically mean, median, standard deviation, and dwell time.

### Calculation of Region Intersection

The intersection of regions is a more complex issue compared to the intersection at the site level. This is because, for region sets *A* and *B*, the number of elements in *A* intersecting *B* is not the same as the number of elements in *B* intersecting *A* . To describe the intersection more accurately, we use the smaller set number between *A* ∩ *B* and *B* ∩ *A*, and the formula is as below,

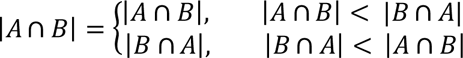

### Gene function enrichment

Gene Ontology (GO) pathways of *E.coli* K12 was obtained online (https://ftp.ebi.ac.uk/pub/databases/GO/goa/proteomes/18.E_coli_MG1655.goa). The GO annotations were obtained with GO.db v3.20.0. The GO enrichment analysis was done by compareCluster v4.8.3 (60) in R v4.3.1.

## Code availability

The custom scripts and plot scripts used for this paper are available in the following GitHub repository: (https://github.com/JeremyQuo/Ecoli_004_ONT_DRS_scripts) and also uploaded to zenodo (https://doi.org/10.5281/zenodo.11664595). And our new method nanoSundial designed for detecting RNA modifications is now available on GitHub at (https://github.com/lrslab/nanoSundial)

## Acknowledgements

We thank the High-Performance Computing Cluster at the City University of Hong Kong for providing us with computational resources. We also thank Hasindu Gamaarachchi from the Garvan Institute of Medical Research for their valuable assistance on GitHub (https://github.com/hasindu2008/f5c/issues/163). Additionally, we are grateful to Ms. Qingqiu Jiang for designing the logo for nanoSundial.

## Contributions

Z.G and R.L designed the research. Y.S and L.T conducted the experiments. Z.G conducted data analysis, visualization and software development. Y.S. conducted the software test. Z.G and Y.S wrote the initial manuscript draft, L.T, B.L, X.D and R.L made corrections and edits. All authors have read and approved the final version of the manuscript.

