## Supplementary figures and tables for "Enhanced Detection of RNA Modifications in *Escherichia coli* Utilizing Nanopore RNA004 Technology": supplementary_file.pdf

### Supplementary materials

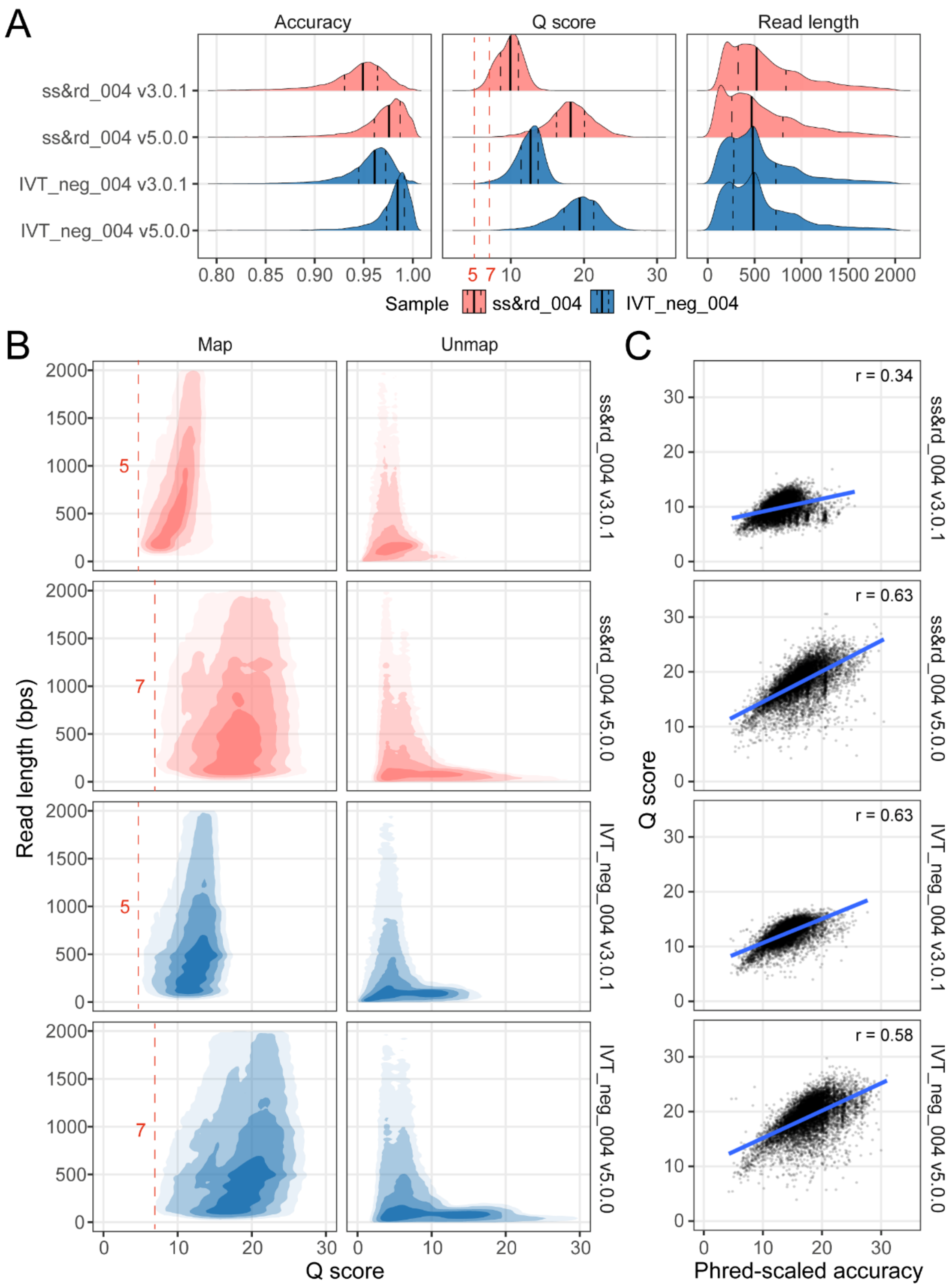


**Figure S1.** Raw read features comparison between different Dorado basecalling model. **(A)** Mapping accuracy, Q scores, and read lengths of mapped reads across different basecall model versions. A Q score of 5 is an effective filter for version 3.0.1, while a Q score of 7 is optimal for version 5.0.0. **(B)** Relationship between read length and the quality of mapped and unmapped reads, presented as a 2D density plot. Read quality is indicated by Q scores. **(C)** Correlation between Q scores and Phred-scaled mapping accuracy across different samples and model versions.

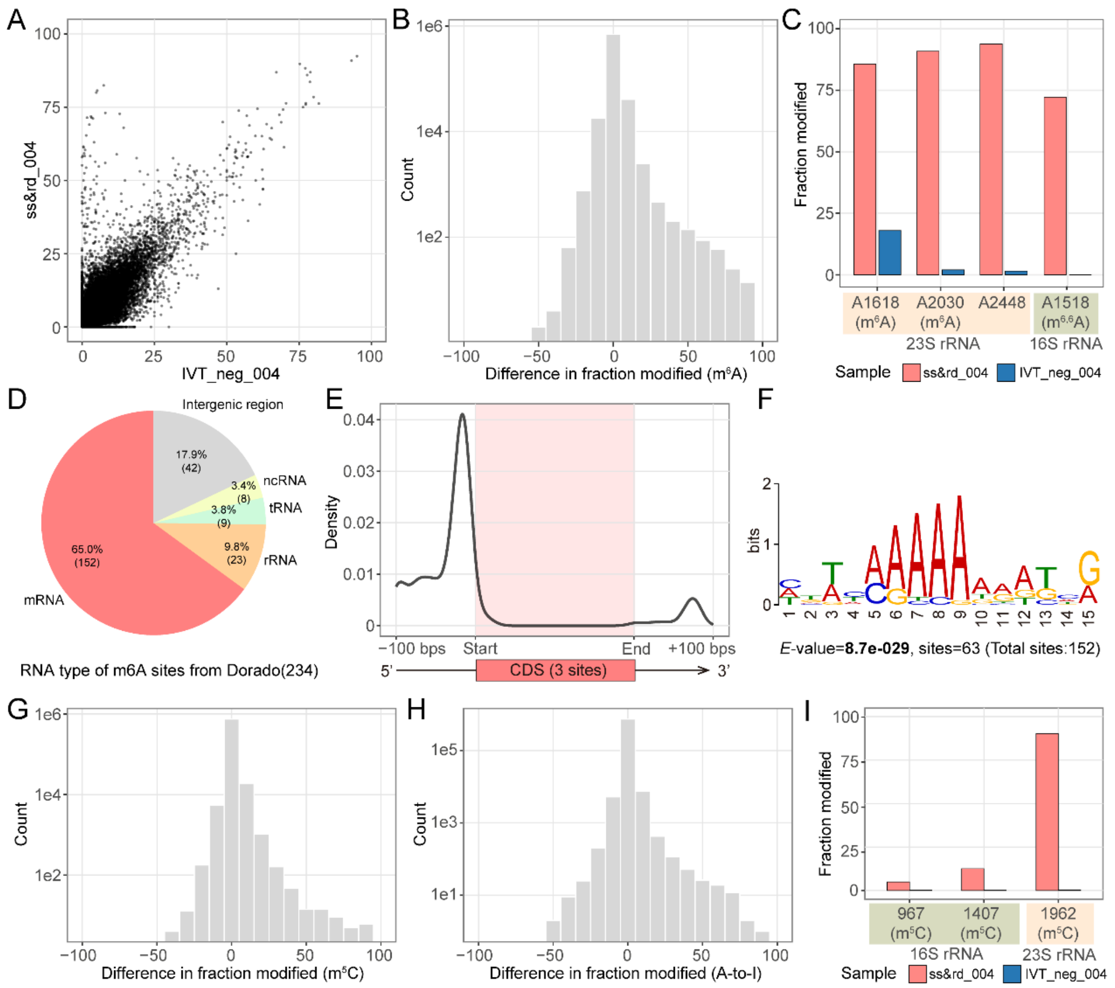


**Figure S2.** Evaluation and characterization of Dorado modification detection model. **(A)** The dot plot illustrates the "fraction modified" values for each position detected by the m6A all-context model across both samples. **(B)** The right-skewed distribution of the difference in "fraction modified" (WT - IVT) is presented, with the y-axis displayed in log10 scale. **(C)** The "fraction modified" values exceeding 65 for E. coli rRNA are highlighted, including two known m6A sites on 23S rRNA (A1618, A2030). After applying a filter of 65, **(D)** Pie plot shows the proportion and number of high-confidence sites within the expanded annotation region. **(E)** The density distribution around the CDS region reveals three sites located within the CDS. **(F)** Motif enrichment analysis using MEME indicates a notable enrichment primarily at the 5' end of the UTR polyA region. **(G)** and **(H)** present the distribution of the difference in "fraction modified" (WT - IVT) for the m5C model and the A-to-I model, respectively. Finally, **(I)** Bar plots illustrate the performance of the m5C model on three known m5C sites within rRNA.

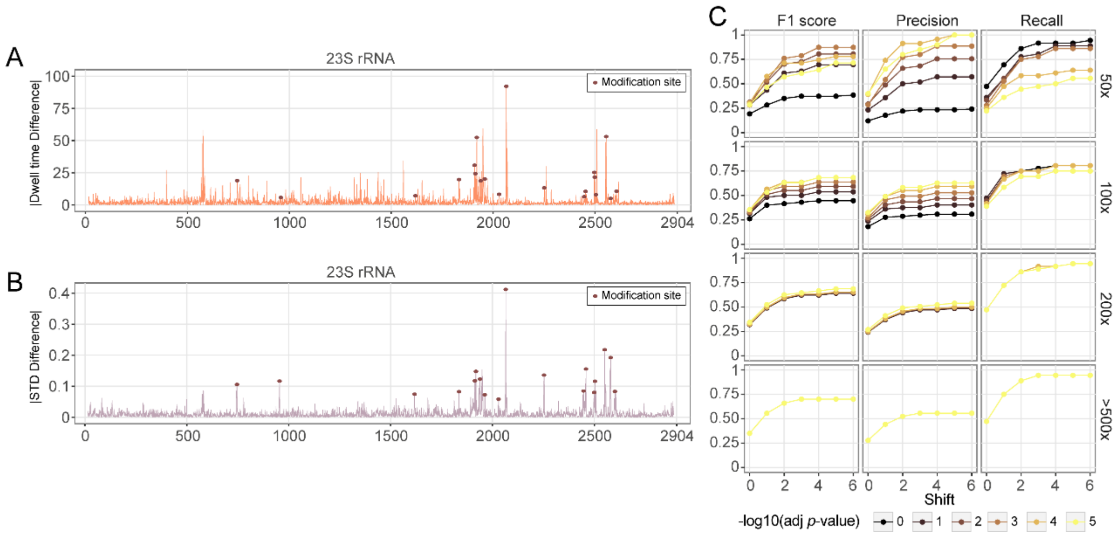


**Figure S3.** Exploration of statistical features of the nanoSundial. **(A)** and **(B)** display the absolute differences in dwell time and standard deviation on 23S rRNA. **(C)** Dot and line plots present the F1 score, precision, and recall values after applying filters based on mean and dwell time. Effects of adjusted *p*-values and shifts were displayed across different coverage levels. Once the coverage reaches a certain threshold (>200x), the *p*-value cutoff becomes ineffective.

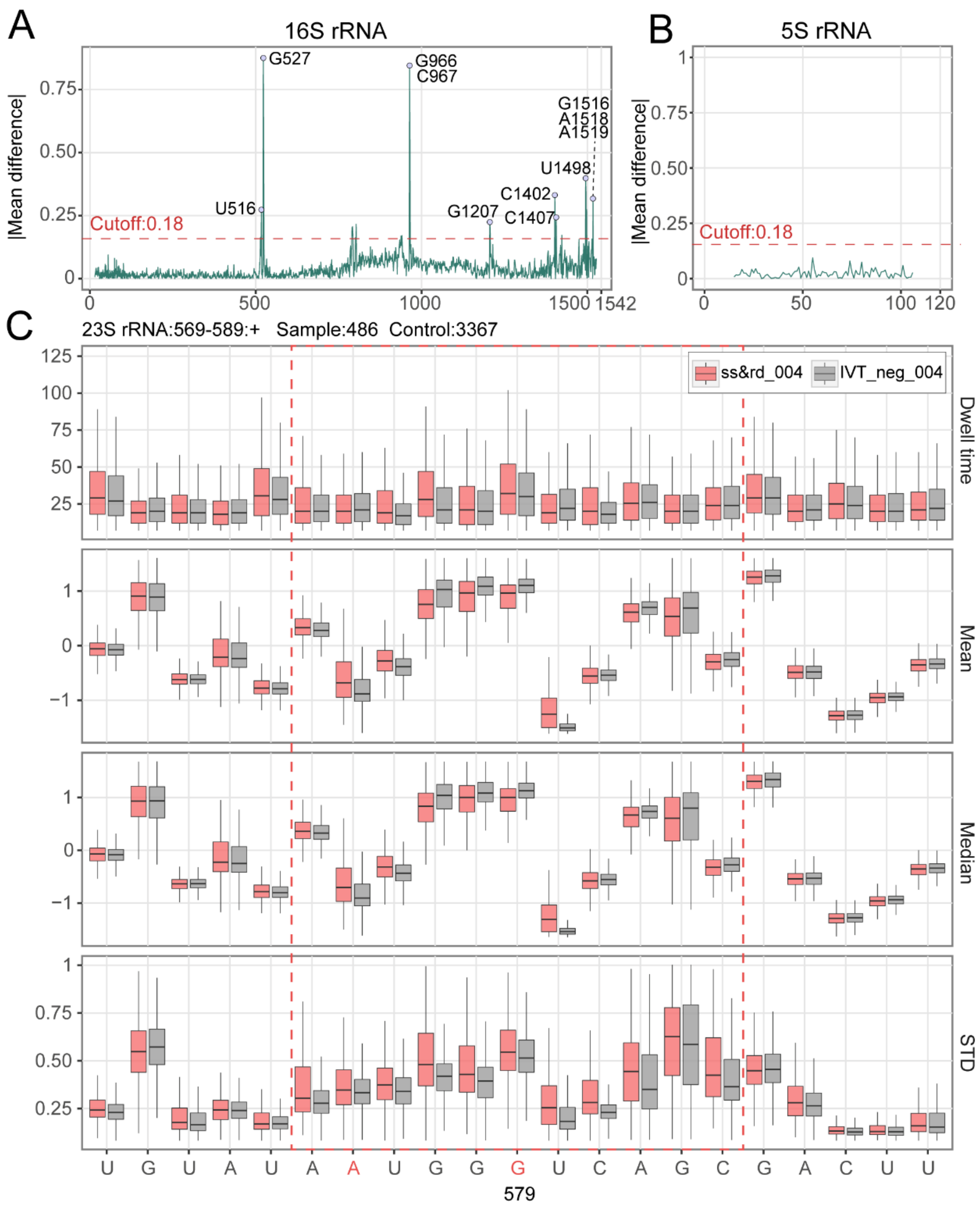


**Figure S4.** Showcase on rRNAs of nanoSundial. **(A)** and **(B)** represent the absolute differences in mean values for 16S and 5S rRNA. **(C)** Box plots illustrate the four features surrounding position 579 on the 23S rRNA, as showcased by nanoCEM.

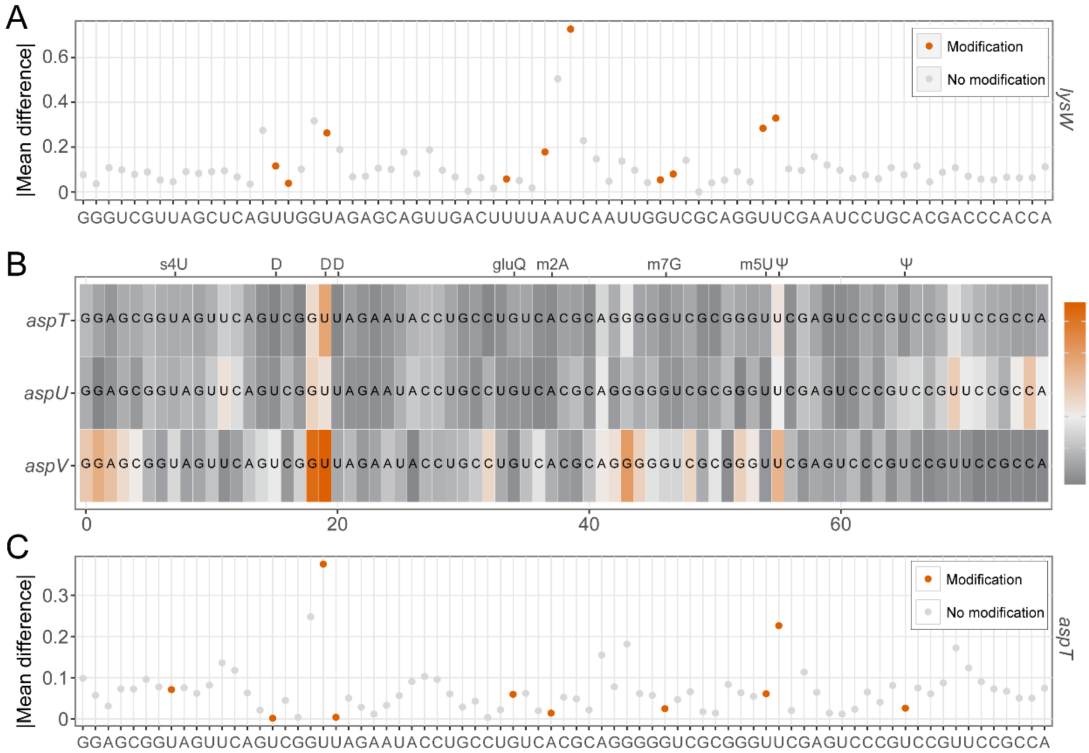


**Figure S5.** Showcase on tRNAs of nanoSundial. **(A)** showcased the absolute value of mean difference on gene *lysW*. **(B)** Heat maps illustrate the absolute value of mean difference for the paralogs of tRNA aspartate (asp). The colors at each position represent the magnitude of these absolute differences. **(C)** A showcase of the absolute value of mean difference on gene *aspT*.

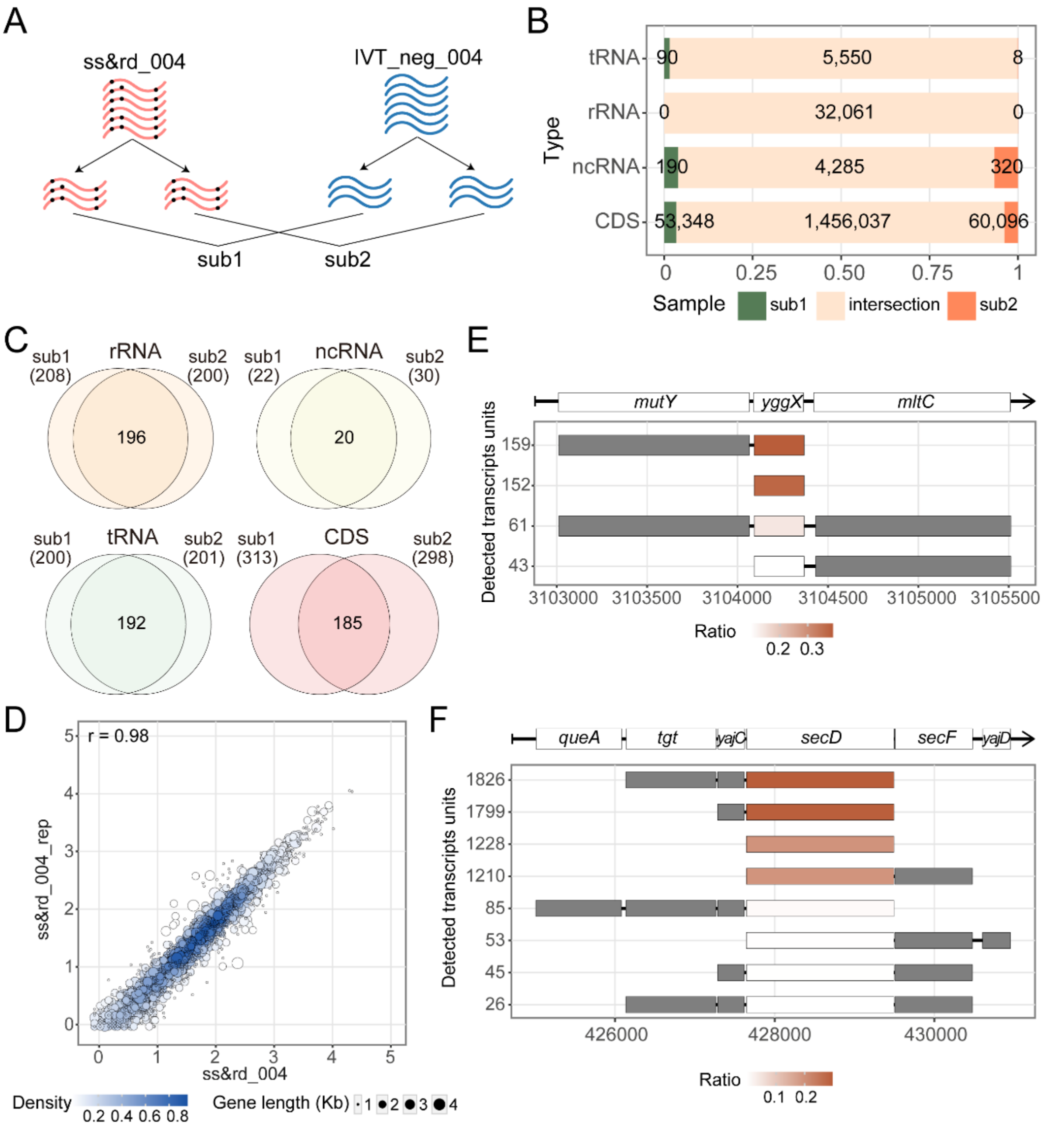


**Figure S6.** Analyzing subsampling and transcription unit structures of stably modified genes **(A)** A workflow to subsample and analyze by nanoSundial. The datasets ss&rd_004 and IVT_neg_004 were divided into two completely non-overlapping subsets, which are then compared using nanoSundial to obtain sub1 and sub2. **(B)** The intersection of the detected sites in sub1 and sub2 within the annotated regions. The green areas represent sites detected only in sub1, the orange areas represent sites detected only in sub2, and the overlapping section indicates sites detected in both sub1 and sub2. **(C)** The intersection of the positive regions from the nanoSundial results for sub1 and sub2. **(D)** Correlations of protein-coding gene expression levels between the ss&rd_004 with respective replicate. Each point represents a single gene, color-coded by density at the plot position. The size of each point indicates gene length. **(E)** and **(F)** The composition and proportion of the Transcription Units (TUs) for *yggX* and *secD*. The colors indicate the proportion of each TU relative to the total reads for one gene.

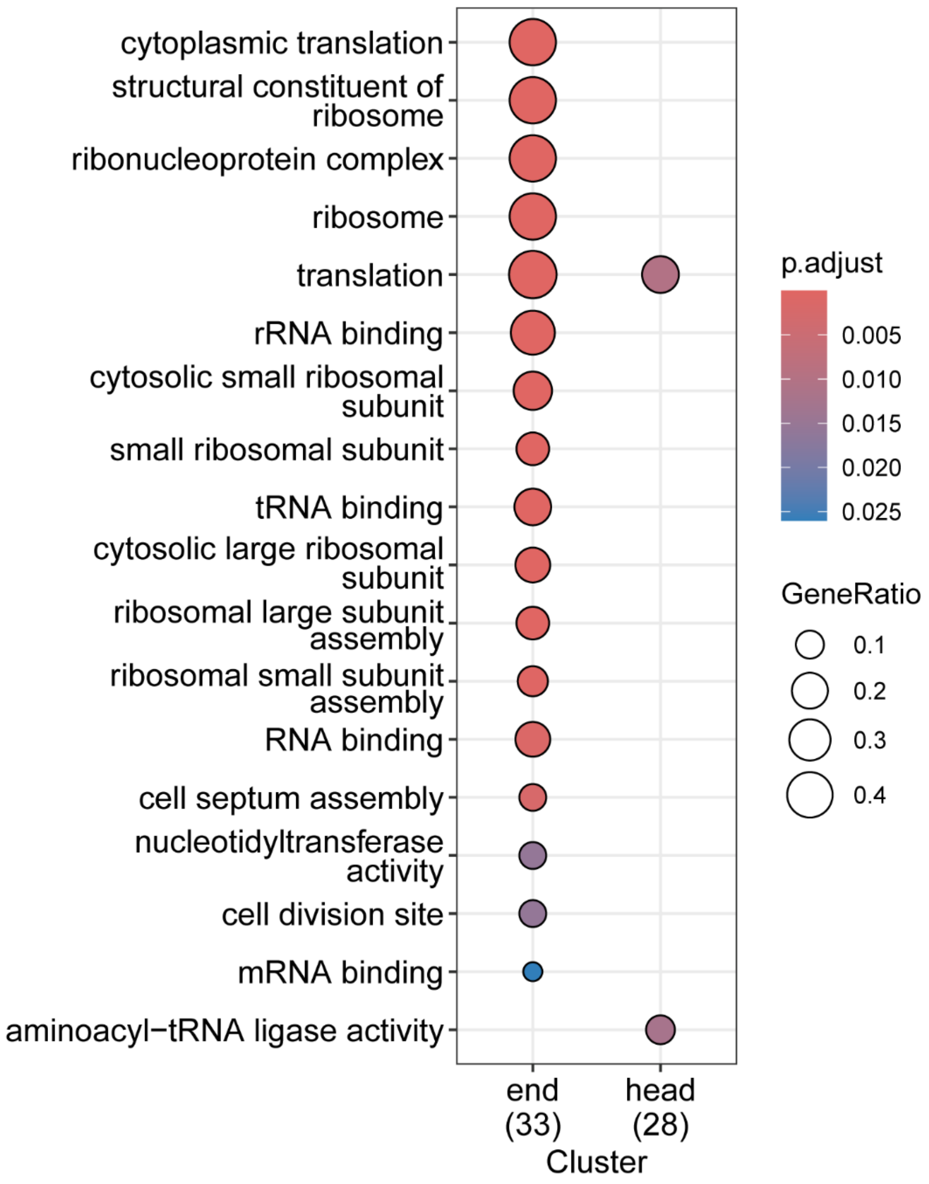


**Figure S7.** GO pathways enrichment of the stably modified genes according to the position in the operons, namely the start or end of the operon.

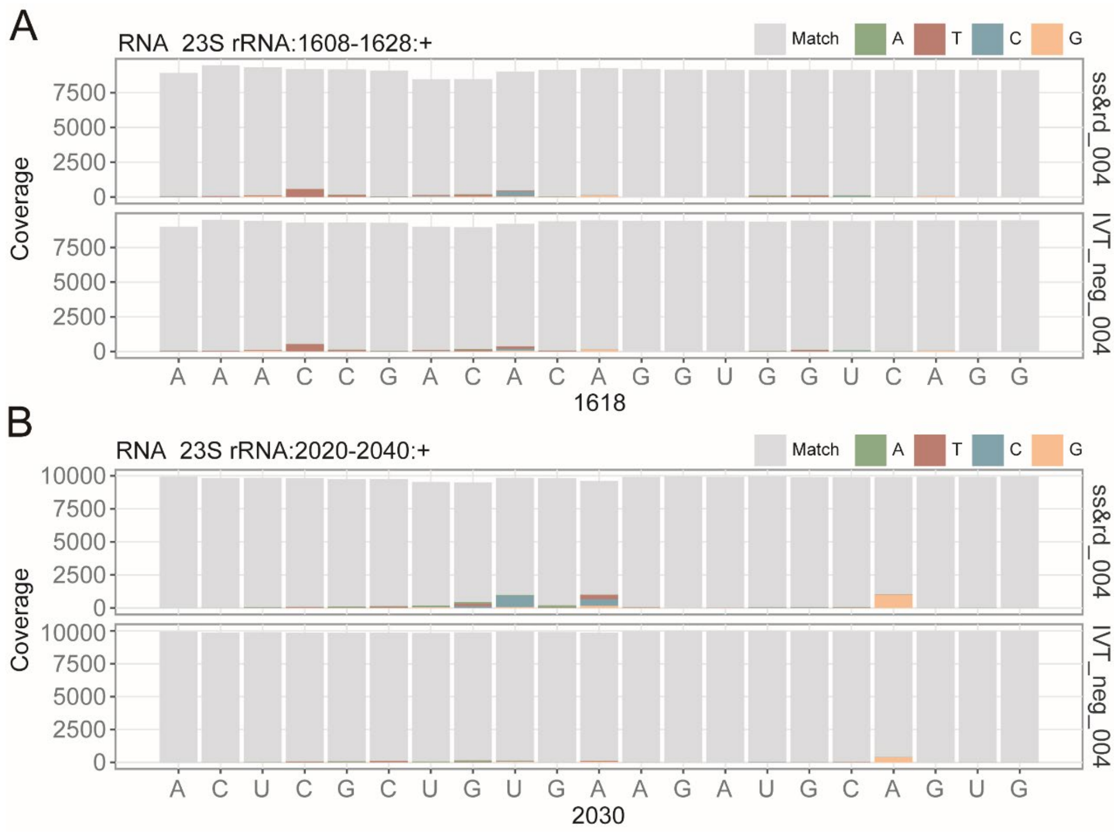


**Figure S8.** Mismatch differences decreased between WT and IVT in RNA004 data**. (A)** and **(B)** illustrate the matches and mismatches of A1618 and A2030, with the gray areas indicating the matching regions of each site.

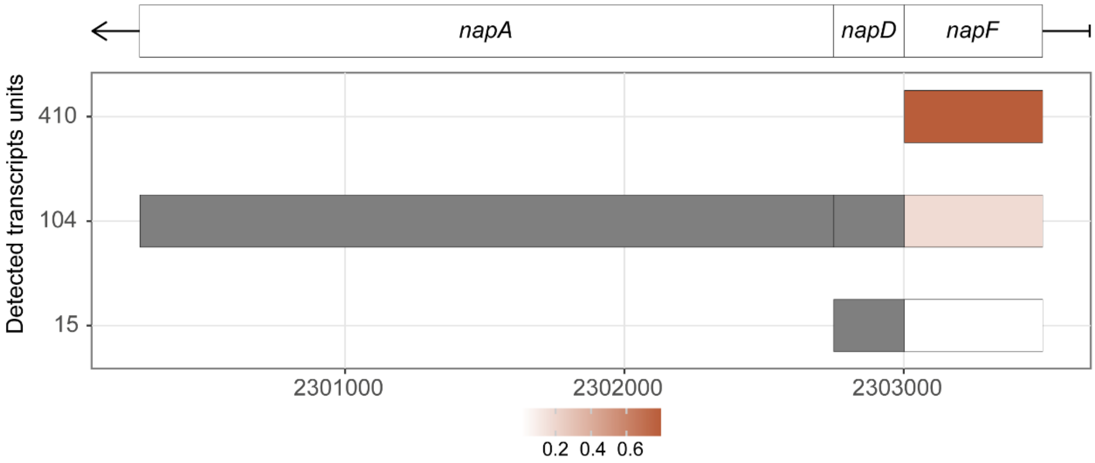


**Figure S9.** The composition and proportion of the Transcription Units (TUs) for *napF*. The colors indicate the proportion of each TU relative to the total reads for one gene.

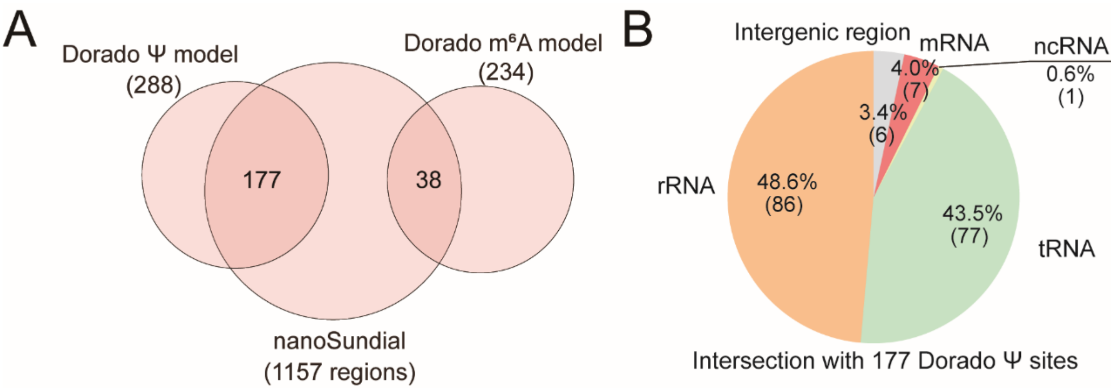


**Figure S10.** The overlap of RNA modifications between Dorado and nanoSundial. **(A)** Overlap between positive regions identified by the Dorado model and nanoSundial. **(B)** RNA type distribution for the 177 sites jointly detected by the Dorado Ψ model and nanoSundial result.

**Table S1**. Sequencing information of raw sequencing data of ss&rd_004 and IVT_004 and their replicates.

**Table S2**. List of high-confidence sites utilizing the "fraction modified" filter from the Dorado Ψ, m6A, m5C, and A-to-I all-context models.

**Table S3**. 714 intersections of positive regions from nanoSundial result of rep1 and pre2, along with the operon information of 71 stably modified gene.
